# Phased chromosome-level genome assembly provides insight into the origin of hexaploid sweetpotato

**DOI:** 10.1101/2024.08.17.608395

**Authors:** Shan Wu, Honghe Sun, John P. Hamilton, Marcelo Mollinari, Gabriel De Siqueira Gesteira, Mercy Kitavi, Mengxiao Yan, Hongxia Wang, Jun Yang, G. Craig Yencho, C. Robin Buell, Zhangjun Fei

## Abstract

The hexaploid sweetpotato (*Ipomoea batatas* [L.] Lam.) is a globally important stable crop and plays a significant role in biofortification. The high resilience and adaptability of sweetpotato provide it with advantages in addressing food security and climate change issues. Here we report a haplotype-resolved chromosome-level genome assembly of an African cultivar, ‘Tanzania’, which enables ancestry inference along the haplotype-phased chromosomes. Our analyses reveal that the wild tetraploid *I. aequatoriensis*, currently found in coastal Ecuador, is the closest known relative of sweetpotato and likely a direct descendant of one of the sweetpotato progenitors. The other unknown progenitor(s) of sweetpotato have a closer genetic relationship to the wild tetraploid *I. batatas* 4×, distributed in Central America, than to *I. aequatoriensis*. The different ancestral sequences are not distributed in typical subgenomes but are intertwined on the same chromosomes, possibly due to the known non-preferential recombination among haplotypes. Although *I. batatas* 4× was not involved in the hexaploidization event, introgression from *I. batatas* 4× to the hexaploid sweetpotato is evident. Our study improves our understanding of sweetpotato origin and provides valuable genomic resources to accelerate sweetpotato breeding.

## Introduction

Sweetpotato (*Ipomoea batatas* [L.] Lam.) is a hexaploid crop (2*n* = 6*x* = 90) originating in South America (Muñoz-Rodríguez et al., 2022) and has been domesticated for more than 5,000 years (Austin, 1988). It was brought to Europe and Africa from Central/South America in the early 16^th^ century and subsequently spread worldwide (Roullier et al., 2013). Sweetpotato is a globally important food security crop with high yield, high nutritional value and adaptability to diverse environmental conditions (Sapakhova et al., 2023). It plays a critical role in the biofortification of provitamin A in sub-Saharan Africa (SSA) to reduce disease caused by vitamin A deficiency. Additionally, sweetpotato provides a vital source of income for subsistence farmers in SSA, who rely on agriculture for their livelihoods. A large number of sweetpotato landraces are grown by SSA sweetpotato farmers, many of which have low yield and are susceptible to diseases (Bashaasha et al., 1995). Robust reference genomes from diploid relatives have advanced sweetpotato breeding and biological research (Wu et al., 2018; Lau et al., 2018; Bednarek et al., 2021; Kitavi et al., 2023). However, hexaploid genomic resources that capture the allelic diversity in sweetpotato are essential for further investigation of its genetics and biology and for more efficient improvement of this important crop.

The origin of sweetpotato remains controversial due to the complexity of its hexaploid genome. Sweetpotato exhibits multivalent chromosomal associations and hexasomic inheritance (Magoon et al., 1970; Mollinari et al., 2020), suggesting that it differs from typical allopolyploids such as the allohexaploid bread wheat (*Triticum aestivum*), which has three clearly distinguishable subgenomes (International Wheat Genome Sequencing Consortium, 2018). An early theory, based on morphological data, suggested the involvement of two diploid relatives, *I. trifida* (2*n* = 2*x* = 30) and *I. triloba* (2*n* = 2*x* = 30) (Austin, 1988), in the formation of hexaploid sweetpotato. Another hypothesis invokes autopolyploidization within *I. trifida* (Roullier et al., 2013; Muñoz-Rodríguez et al., 2018). The discovery of a wild autotetraploid species, *I. aequatoriensis* (2*n* = 4*x* = 60), closely related to sweetpotato, has led to a third hypothesis involving allopolyploidization between *I. trifida* and *I. aequatoriensis* (Muñoz-Rodríguez et al., 2022). The most recent hypothesis suggests the contribution of wild tetraploid *I. batatas* (*I. batatas* 4×; 2*n* = 4*x* = 60) and *I. aequatoriensis* to the hexaploid sweetpotato genome (Yan et al., 2024). The absence of a phased hexaploid genome assembly has been a major barrier to understanding the origin and evolution of sweetpotato.

In this study, we generated a fully phased 90-chromosome assembly of an African sweetpotato cultivar, ‘Tanzania’, which is widely grown in SSA. By analyzing sequence similarity and genetic relationships between sweetpotato haplotypes and the existing closely related wild species, we determined the involvement of at least two species in the hexaploidization event, one of which could have directly descended into the wild *I. aequatoriensis*, currently restricted to coastal Ecuador, while the other genome donor(s) remain unknown but are closely related to the wild Central American *I. batatas* 4×. Ancestry inference along the chromosomes revealed extensive exchange and replacement of different ancestral sequences among the six haplotypes.

This study enhances our understanding of sweetpotato origin and evolution and provides a valuable genomic resource to facilitate sweetpotato breeding and trait research.

## Results

### Genome of the hexaploid sweetpotato ‘Tanzania’

To explore the complexity and origin of the hexaploid sweetpotato genome, we developed a chromosome-level haplotype-resolved assembly for ‘Tanzania’ (**Fig. 1a**), the most widely grown sweetpotato cultivar in SSA, featuring high storage root dry matter content and resistance to sweetpotato virus disease and *Alternaria* stem blight (Mwanga et al., 2001). ‘Tanzania’ has an estimated genome size of approximately 2.78 Gb according to k-mer analysis (**Supplementary Fig. 1**). A total of 128.75 Gb of PacBio HiFi and 563.67 Gb of chromatin conformation capture (Hi-C) sequencing data were generated, representing approximately 43× and 188× coverage of the hexaploid ‘Tanzania’ genome, respectively. The final phased assembly was 2.76 Gb with a contig N50 length of 3.77 Mb (**Supplementary Table 1**). The assembly size was close to the estimated hexaploid genome size, and a single-peak HiFi read depth distribution pattern across the assembly was observed (**Supplementary Fig. 2**), indicating minimal collapse of highly duplicated regions in the assembly. Consistently, k-mer spectrum analysis indicated that the assembly captured most of the heterozygous content in the hexaploid genome (**Supplementary Fig. 3**). Furthermore, k-mer analysis using Merqury (Rhie et al., 2020) indicated the high base accuracy and completeness of the assembly, with a consensus quality value (QV) of 61.29 and a k-mer completeness rate of 98.92%. The ‘Tanzania’ genome assembly also exhibited a high LTR assembly index (Ou et al., 2018) of 11.34. The completeness of the assembly was further assessed using BUSCO (Simao et al., 2015), revealing that 99.5% of the core conserved plant genes were completely captured in the assembly, with 98.5% duplicated and 64.9% present in six copies (**Supplementary Table 2** and **Supplementary Fig. 4**).

**Fig. 1.**
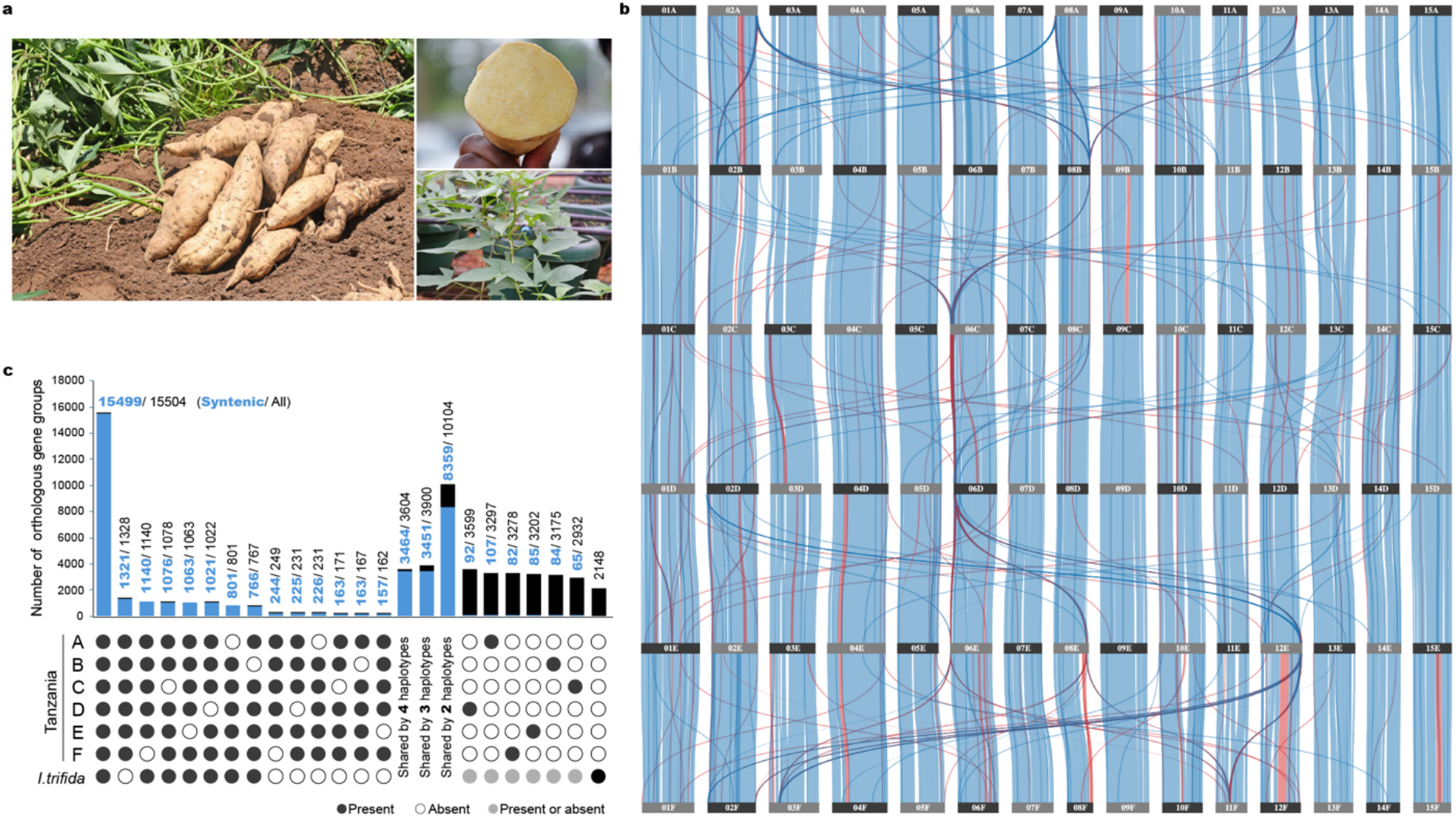
Phased genome assembly of the hexaploid sweetpotato cultivar ‘Tanzania’. **a**, Images of ‘Tanzania’ storage roots and seedlings. **b**, Synteny map among the six haplotypes of ‘Tanzania’ (haplotype A to F from top to bottom). The synteny map was generated using Synvisio (https://synvisio.github.io). Alignment blocks with forward and reverse orientations are labeled in blue and red, respectively. **c**, Orthologous and paralogous gene groups identified in the six sweetpotato haplotypes and *I. trifida* NCNSP0306.

For haplotype phasing and construction of a chromosome-level assembly of the ‘Tanzania’ genome, we used a combination of evidence, including phased genetic maps constructed from a mapping population with ‘Tanzania’ being one of the parents (Mollinari et al., 2020), Hi-C contact signals, and synteny to the 15-chromosome consensus assembly that was constructed by anchoring the primary unphased contigs into pseudochromosomes using genetic markers (Mollinari et al., 2020). The consensus assembly had a total length of 455.6 Mb, close to the size of a monoploid genome, and a contig N50 size of 13.4 Mb (**Supplementary Tables 1-3**). Simplex markers from the hexaploid sweetpotato genetic map with fully phased haplotypes for ‘Tanzania’ (Mollinari et al., 2020) assigned 822 contigs with a total length of 2.50 Gb (90.39% of the assembly) to 90 haplotype groups. Among these contigs, 44 were found to be chimeric and were broken at phase switch sites indicated by genetic markers and depleted Hi-C contact signals (**Supplementary Fig. 5**). Hi-C contact signals between contigs were then used to assign the remaining contigs to the 90 haplotypes. The pseudochromosomes were constructed within each haplotype, resulting in anchoring and orientating 94.92% (2.62 Gb) of the phased assembly into 90 pseudochromosomes (**Supplementary Table 3**). The correctness of phasing and scaffolding at the chromosome level was supported by Hi-C contact maps (**Supplementary Figs. 6**). The phasing accuracy was further evaluated using nPhase (Abou Saada et al., 2021), which revealed a very low level of potential phase switching errors (0.49%) in the ‘Tanzania’ genome assembly (**Supplementary Fig. 7**). Moreover, we constructed a phased ‘Tanzania’ genetic map using the Beauregard × Tanzania population described in Mollinari et al. (2020), with the phased ‘Tanzania’ genome assembly as the reference. Comparison of the phased genetic map and the phased genome assembly revealed a consistency rate of 98.6% (**Supplementary Fig. 8**). Together, these results confirmed the high accuracy of the phased chromosome-level assembly of the ‘Tanzania’ genome.

Genome alignment revealed strong collinearity among the six haplotypes, despite the presence of seven large inversions with sizes ranging from 737 kb to 5.66 Mb (**Fig. 1b, Supplementary Figs. 9** and **10** and **Supplementary Table 4**). These inversions were supported by Hi-C contact signals and HiFi read alignments (**Supplementary Fig. 11**). Additionally, 17,313,423 biallelic SNPs were identified among the six haplotypes. Of the 10,203,166 SNP sites with all six haplotypes genotyped, 5,663,151 (55.5%) exhibited a 1:5 allele ratio (simplex), 3,280,996 (32.2%) had a 2:4 ratio (duplex), and 1,259,019 (12.3%) showed a 3:3 (triplex) ratio.

The predominance of simplex sites was consistent with previous allele dosage analysis in 16 sweetpotato cultivars (Wu et al., 2018). Insertions and deletions (indels) larger than 20 bp were also identified among the six haplotypes (**Supplementary Table 5**). Using haplotype A as the reference, 285,118 non-redundant insertions (total length: 230.3 Mb) from the other five haplotypes were detected, and 176,125 sequences with a total length of 113.8 Mb were present in haplotype A but absent in at least one other haplotype. More than half of these indels (254,009 out of 461,243) were smaller than 100 bp, while indels ranging from one to ten kb contributed to more than half of the total inserted and deleted bases among the haplotypes (**Supplementary Table 5**).

### Gene retention and expansion in the hexaploid sweetpotato genome

Approximately 55.7% of the ‘Tanzania’ assembly was identified as repetitive sequences (**Supplementary Table 6**). A total of 246,023 high-confidence protein-coding genes were predicted from the ‘Tanzania’ genome, capturing 98.6% of the conserved BUSCO genes (**Supplementary Table 2**). Of these, 231,798 genes (94.22%) were located on the 90 chromosomes, with an average of 38,633 genes per haplotype. About 79.54% (195,692) of the predicted genes had hits to functional domains in the InterPro database, and 85.84% (211,196) to the GenBank non-redundant (nr) protein database. A total of 39,654 syntenic homoeologous gene groups were identified among the six haplotypes of ‘Tanzania’ and the wild relative, *I. trifida* ‘NCNSP0306’ (Wu et al., 2018), comprising 202,439 *I. batatas* genes and 27,712 *I. trifida* genes. Among these groups, 23,865 (155,824 genes) were shared by at least five haplotypes, and 16,820 were shared by all six haplotypes, with 13,782 having a 1:1:1:1:1:1 ratio (**Fig. 1c** and **Supplementary Table 7**).

Although most sweetpotato genes were retained in multiple homoeologous copies following the polyploidization event, 22,100 genes were present in only one haplotype (**Fig. 1c**). Housekeeping genes performing core functions and highly conserved across eukaryotes are often not retained as duplicates after genome duplications in Angiosperms (De Smet et al., 2013). Indeed, functions of these haplotype-specific genes were enriched in DNA integration, DNA biosynthesis, and biological processes related to cytoskeleton-dependent trafficking (**Supplementary Table 8**), suggesting that retaining multiple copies of such housekeeping genes might not be beneficial to the fitness of hexaploid sweetpotato. In contrast, genes with functions related to stress response and disease resistance tend to be retained in multiple copies in polyploid species, either due to human selection or their roles in environmental adaptation (Geiser et al., 2016; IWGSC, 2018). Functional gene families were identified based on protein domains in the phased hexaploid ‘Tanzania’ (90 chromosomes) and *I. trifida* ‘NCNSP0306’ (15 chromosomes) genomes. For most gene families, their sizes in the hexaploid *I. batatas* genome were six times those in the haploid *I. trifida* genome assembly (**Supplementary Fig. 12**). Gene families with noticeably larger sizes included the TIR-domain family, NBS-ARC, and leucine-rich repeat genes (**Supplementary Fig. 12** and **Supplementary Table 9**). A total of 616 and 53 TIR-NBS-LRR genes were annotated in the phased *I. batatas* ‘Tanzania’ and the haploid *I. trifida* assemblies, respectively. The largest cluster of TIR-NBS-LRR genes in ‘Tanzania’ was found on the F haplotype of chromosome 7 from 24.8-25.8 Mb, containing 60 TIR-NBS-LRR genes (**Supplementary Fig. 13**). Interestingly, 36 out of these 60 genes in this cluster were found in one syntenic homoeologous group, which also contained five other TIR-NBS-LRR genes in haplotype A (five out of a cluster of 26 TIR-NBS-LRR genes) and one TIR-NBS-LRR gene in haplotype E (**Supplementary Fig. 14**). The observed distribution patterns of groups of syntenic homoeologous TIR-NBS-LRR genes on the six haplotypes of chromosome 7 suggest that these large TIR-NBS-LRR gene clusters could have resulted from multiple segmental duplication events of tandemly arrayed TIR-NBS-LRR genes.

### *I. aequatoriensis* directly descended from one donor of the hexaploid sweetpotato

The haplotype-resolved chromosome-level genome assembly of ‘Tanzania’ enabled us to analyze the origin of sweetpotato. Previous studies involving extensive sampling of related wild diploid and tetraploid species have established two tetraploid species, *I. aequatoriensis* and *I. batatas* 4×, as the closest extant wild relatives of cultivated sweetpotato (Muñoz-Rodríguez et al., 2022; Yan et al., 2024). Using genome resequencing data from ten *I. aequatoriensis* and eight *I. batatas* 4× accessions (Yan et al., 2024; Muñoz-Rodríguez et al., 2022), we explored the potential contributions of *I. aequatoriensis* and *I. batatas* 4× to the hexaploid genome. We first classified each ‘Tanzania’ genomic window as either *I. aequatoriensis* or *I. batatas* 4× type based on the genetic distance between ‘Tanzania’ and the two wild species. Phylogenetic trees were constructed using ‘Tanzania’ and the 18 wild accessions across 22,348 non-overlapping ‘Tanzania’ genomic windows, each containing 5,000 SNPs and ranging from 51,379 bp to 3,145,378 bp in size. Ancestry was also inferred based on sequence identity between each wild accession and ‘Tanzania’ in the phased genomic windows. The distribution of sweetpotato-versus-wild sequence identity differences of the two wild species displayed two peaks, corresponding to ‘Tanzania’ genomic regions with higher sequence identities to either *I. aequatoriensis* or *I. batatas* 4× (**Supplementary Fig. 15**). Two normal distributions were fitted to this pattern (**Supplementary Fig. 15**), and the 95% confidence intervals were chosen as the cutoff for determining whether a ‘Tanzania’ region was more similar to *I. aequatoriensis* or *I. batatas* 4×. Both *I. aequatoriensis* and *I. batatas* 4× showed higher sequence identities to the hexaploid sweetpotato than *I. trifida* (**Fig. 2a**), excluding the possibility of *I. trifida* being a direct donor of the sweetpotato genome. The distribution of sequence identities between *I. aequatoriensis* and ‘Tanzania’ showed a peak at over 99.5% identity (**Fig. 2a**). A total of 5,079 ‘Tanzania’ windows (603.3 Mb) had higher sequence identity to *I. aequatoriensis* than to *I. batatas* 4×, of which 4,818 (567.1 Mb; 21.7% of the phased genome) were also classified as *I. aequatoriensis* type based on genetic distance (**Fig. 2b** and **Supplementary Fig. 16**). These high-confidence *I. aequatoriensis*-like regions in ‘Tanzania’ (determined by both sequence identity and genetic distance) were not evenly distributed among the six haplotypes nor concentrated in just one or two haplotypes (**Fig. 2c** and **Supplementary Fig. 16**). The 15 homoeologous chromosome groups also contained varying amounts of *I. aequatoriensis*-like sequences, ranging from 11.2% in chromosome 7 to 37.3% in chromosome 13 (**Supplementary Fig. 16**).

**Fig. 2.**
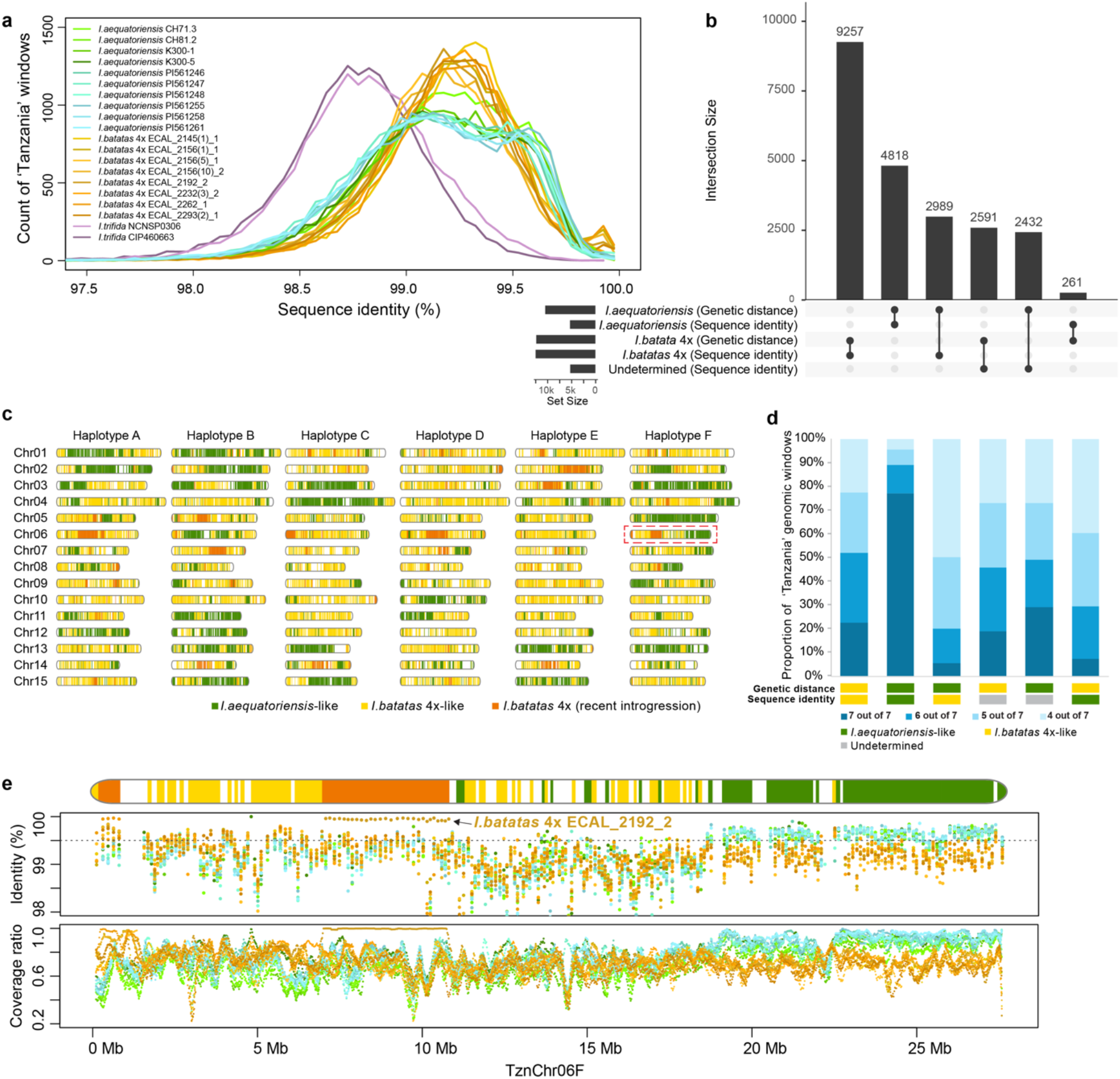
Ancestry inference of hexaploid sweetpotato. **a**, Distribution of sequence identities between ‘Tanzania’ and wild *I. aequatoriensis, I. batatas* 4×, and *I. trifida* accessions. **b**, Numbers of genomic windows inferred to have *I. aequatoriensis* or *I. batatas* 4× ancestry based on genetic distance and sequence identity. **c**, Ideograms of the 90 ‘Tanzania’ chromosomes illustrating local ancestry and introgression. Each vertical line represents a genomic window with 5,000 non-overlapping SNPs. **d**, Proportions of ‘Tanzania’ genomic windows for which ancestry inference based on genetic distance was supported by four to seven nearest neighboring accessions. Each column represents windows with consistent or inconsistent ancestry inference based on genetic distance and sequence identity, ordered as in panel **b. e**, Ancestry inference along the ‘Tanzania’ chromosome 6F (highlighted in **c**). Sequence identities between sweetpotato and wild tetraploid accessions are plotted below the chromosome ideogram. Coverage ratios of the ‘Tanzania’ genome by genomic reads of wild tetraploid accessions in 200-kb windows with a 20-kb step size are shown at the bottom. Wild accessions are labeled with colors corresponding to those in **a**.

On the other hand, the *I. batatas* 4×-like sequences in ‘Tanzania’, supported by both sequence identity and genetic distance, had a total size of 1,104.4 Mb (9,257 windows), approximately twice the size of the *I. aequatoriensis*-like sequences (**Fig. 2b** and **Supplementary Fig. 16**). While the ten *I. aequatoriensis* individuals exhibited a peak sequence identity of 99.6% to the *I. aequatoriensis*-like sequences in the ‘Tanzania’ genome, the eight *I. batatas* 4× individuals showed a lower sequence identity (99.3%) to the identified *I. batatas* 4×-like sequences (**Supplementary Fig. 17**). Given the similar mutation rates of sequences within the same hexaploid sweetpotato genome contributed by different donors, this lower sequence identity suggests that *I. batatas* 4× may not be the direct descendant of the other donor of the hexaploid sweetpotato genome. Furthermore, only 20% of the *I. batatas* 4×-type windows, classified by both sequence identity and genetic distance methods, were confidently supported by all seven individuals with the closest genetic distance being *I. batatas* 4× accessions (**Fig. 2d**). In contrast, 80% of the *I. aequatoriensis*-type windows had all nearest neighbors exclusively from *I. aequatoriensis* (**Fig. 2d**). Taken together, while *I. aequatoriensis* is likely to be the direct descendant of one of the progenitors of the hexaploid sweetpotato, the other direct donor(s) might not have been sampled yet. The unsampled species were likely more closely related to *I. batatas* 4× than to *I. aequatoriensis*, as suggested by the sequence identity.

### *Ib*T-DNAs in the ‘Tanzania’ genome

Horizontally transferred *Ib*T-DNA sequences are ubiquitously present in hexaploid sweetpotato genomes (Kyndt et al., 2015). These *Ib*T-DNAs are also found in wild relatives of sweetpotato and have been used to trace the origins of sweetpotato (Quispe-Huamanquispe et al., 2019; Yan et al., 2024). While *Ib*T-DNA1 is detected in all 291 globally sampled hexaploid sweetpotato accessions, *Ib*T-DNA2 is present in only about 20% of analyzed sweetpotato cultivars (Kyndt et al., 2015). Nearly all *I. aequatoriensis* accessions possess *Ib*T-DNA2, and some *I. batatas* 4× accessions contain *Ib*T-DNA1 (Yan et al., 2024). ‘Tanzania’ has been found to contain both *Ib*T-DNA1 and *Ib*T-DNA2 (Kyndt et al., 2015). In ‘Tanzania’, the *Ib*T-DNA1 sequence was detected at the end of chromosome 14 in haplotypes C, D and E (**Supplementary Fig. 18a**). The two inverted repeats of *Ib*T-DNA1 in ‘Tanzania’ (**Supplementary Fig. 18b**) have previously been described in the sweetpotato variety ‘Xu781’ (Kyndt et al., 2015). *Ib*T-DNA2 insertions were found at the end of chromosome 4 in haplotypes A, B and D, with their inverted repeat patterns also observed in ‘Tanzania’ (**Supplementary Fig. 18**). Interestingly, the *Ib*T-DNA1 and *Ib*T-DNA2 insertions in ‘Tanzania’ were found in genomic regions with *I. batatas* 4×-like and *I. aequatoriensis* ancestry, respectively (**Supplementary Fig. 18a**), consistent with the finding that *Ib*T-DNA1 and *Ib*T-DNA2 are present in current *I. batatas* 4× and *I. aequatoriensis* populations, respectively (Yan et al., 2024). The presence of three copies of *I. batatas* 4×-like *Ib*T-DNA1 and three copies of *I*.

*aequatoriensis*-like *Ib*T-DNA2 in the ‘Tanzania’ genome might seem unexpected, given that our ancestry inference suggested that the *I. batatas* 4×-like progenitor contributed twice as much to the hexaploid genome as *I. aequatoriensis*. However, this could be attributed to gene conversion among randomly paired haplotypes and the mixing of different alleles during hybridization between sweetpotato accessions, leading to various dosages of *Ib*T-DNA1 and *Ib*T-DNA2 in sweetpotato individuals, with complete loss of *Ib*T-DNA2 in many cultivars as an extreme case (Kyndt et al., 2015).

### Wild *I. batatas* 4× introgression in the cultivated sweetpotato

Although *I. batatas* 4× was not directly involved in the hexaploidization event of sweetpotato, introgressions from extant *I. batatas* 4× into the hexaploid sweetpotato were observed (**Fig. 2c**). A total of 1,031 genomic windows in ‘Tanzania’, with a total length of 144.3 Mb, were identified as potentially introgressed from *I. batatas* 4× (**Supplementary Table 10**). Notably, the genomic region on chromosome 6F of ‘Tanzania’, from 7.0 to 10.8 Mb, exhibited nearly identical sequences to the *I. batatas* 4× accession ECAL_2192_2 (**Fig. 2e**). The 8,662 genes within these introgressed regions were enriched with those involved in iron-sulfur (Fe-S) cluster assembly and meiotic chromosome segregation (**Supplementary Table 11**). Fe-S clusters play key roles in plant development, including photosynthesis, respiration, nitrogen assimilation, and disease resistance (Balk and Pilon, 2011; Fonseca et al., 2020). Meiotic chromosome segregation is crucial for ensuring balanced gametes and preventing aneuploidy in polyploid plants (Comai, 2005). Taken together, the introgression of wild *I. batatas* 4× sequences into sweetpotato might support essential developmental processes and overcome challenges associated with polyploid meiosis. The contribution of different *I. batatas* 4× accessions to the introgressions varied, ranging from 3.3 Mb in ECAL_2156(5)_1 to 46.7 Mb in ECAL_2262_1. Additionally, approximately 78.3 Mb of the introgressed sequences (54.3% of the total) were detected in only one *I. batatas* 4× accession. For example, 32.6 Mb of introgressed sequences (22.6% of the total) could only be found in the accession ECAL_2262_1 (**Supplementary Fig. 19**).

## Discussion

The origin of hexaploid sweetpotato has been a topic of debate for decades. The discovery of the new Ecuadorian species *I. aequatoriensis* and recent phylogenetic analyses have shed light on potential ancestors of sweetpotato (Muñoz-Rodríguez et al., 2022; Yan et al., 2024). With our phased chromosome-level hexaploid sweetpotato genome assembly, we can now test various hypotheses about sweetpotato origin based on actual sequence similarities and phylogenetic relationships among the six sweetpotato haplotypes and closely related wild species. Our analyses support the allopolyploid hypothesis for sweetpotato origin. Approximately 63.8% of the phased ‘Tanzania’ genome could be confidently inferred to have an *I. aequatoriensis* or *I. batatas* 4×-like ancestry, with about one-third contributed by an *I. aequatoriensis*-related progenitor. The wild tetraploid *I. aequatoriensis* shows the highest sequence similarity to the hexaploid sweetpotato, suggesting that it could be a direct descendant of one progenitor of sweetpotato. This also implies that sweetpotato could originate in tropical South America, where *I. aequatoriensis* is currently distributed (Muñoz-Rodríguez et al., 2022). While some chromosomes in the hexaploid sweetpotato ‘Tanzania’ genome, such as chromosomes 3F, 4C, 5F and 11B, were predominantly occupied by the *I. aequatoriensis*-like sequences (**Fig. 2c**), an intertwined pattern of *I. aequatoriensis*- and *I. batatas* 4×-like sequences along the chromosomes is prevalent. This pattern could result from the known autopolyploid-like random pairing among homoeologous chromosomes during meiosis in hexaploid sweetpotato (Mollinari et al., 2020), allowing for the exchange and replacement of sequences initially donated by different progenitors. The much lower sequence identity between the extant diploid *I. trifida* and hexaploid sweetpotato than that between *I. aequatoriensis* or *I. batatas* 4× and sweetpotato disclaims the direct involvement of *I. trifida* in the hexaploidization event, which is also supported by previous chloroplast genome analyses (Yan et al., 2024). The relatively low sequence similarity between *I. batatas* 4× and sweetpotato suggests that the unsampled direct donor(s) of the sweetpotato genome and *I. batatas* 4× likely share a common ancestor that diverged earlier from *I. aequatoriensis*. Therefore, sequences in the hexaploid sweetpotato genome contributed by the unknown donor(s) have a closer genetic relationship to *I. batatas* 4× than to *I. aequatoriensis*. Notably, a small portion (about 5%) of the sweetpotato genome was nearly identical to *I. batatas* 4×, suggesting potential introgressions from wild *I. batatas* 4× into hexaploid sweetpotato.

The concept of “segmental” allopolyploid was introduced by Stebbins (1947) to describe allopolyploids that arise from the genome merger of divergent donors and exhibit multivalent cytological behavior, resembling autopolyploids during meiosis more than true allopolyploids. In the case of hexaploid sweetpotato, which has genetically distinct genome donors that are potentially from different species, it be classified as an allopolyploid by definition. However, the sweetpotato genome lacks mechanisms to strictly discriminate subgenomes and enforce bivalent homologous pairing. Instead, it exhibits non-preferential and multivalent pairing of homoeologous chromosomes, as supported by cytological and genetic evidence (Magoon et al., 1970; Mollinari et al., 2020), which has allowed the shuffling of donor sequences, leading to haplotype blocks with mixed sequences from different progenitors. Given the divergent donors and autopolyploid-like hexasomic inheritance, it is reasonable to tentatively classify sweetpotato as a segmental allopolyploid.

Diploid reference genomes have been fundamental for sweetpotato breeding and functional studies (Wu et al., 2018; Lau et al., 2018; Bednarek et al., 2021; Kitavi et al., 2023). However, they cannot fully capture the genetic and allelic diversity present in hexaploid sweetpotato. The haplotype-resolved chromosome-level genome assembly of ‘Tanzania’ represents a significant milestone in genomics-assisted sweetpotato improvement. This assembly allows for a detailed exploration of the complex sweetpotato genome, enhances our understanding of its origin and evolution, and improves our ability to study genetic controls of agronomic traits with respect to different functional alleles and dosage effects. During evolution and breeding, alleles may be lost in genomes of single sweetpotato cultivars due to gene conversion and various allele combinations. For example, *I. aequatoriensis*-like alleles on ‘Tanzania’ chromosome 7 were largely replaced by alleles from other progenitor(s) (**Fig. 2c** and **Supplementary Fig. 16**), and many sweetpotato accessions may have lost the allelic regions carrying *Ib*T-DNA2 (Kyndt et al., 2015). Generating additional haplotype-resolved genome assemblies of sweetpotato cultivars from diverse breeding programs will provide a more comprehensive view of genomic variations and allele inheritance. This will facilitate the development of more efficient strategies for sweetpotato improvement.

## Methods

### Genome sequencing

High molecular weight genomic DNA of ‘Tanzania’ was extracted from fresh leaves using the cetyltrimethylammonium bromide (CTAB) method and used for the construction of a SMRTbell library using the SMRTbell Express Template Prep Kit 2.0 (PacBio). The library was sequenced with the circular consensus sequencing (CCS) mode on a PacBio Sequel II instrument to generate HiFi reads. A Hi-C library was prepared following the proximo Hi-C plant protocol (Phase Genomics) and sequenced on an Illumina NextSeq 500 platform with the paired-end mode and a read length of 150 bp.

### Phased genome assembly and pseudochromosome construction

The haplotype-resolved contigs of ‘Tanzania’ were assembled from PacBio HiFi and Hi-C sequencing data using hifiasm (v0.19.5-r587) (Cheng et al., 2022). Potential contaminations were detected by aligning the contigs to the NCBI nonredundant nucleotide (nt) database using BLASTN with an E value cutoff of 1e-5. Contigs with more than 80% of their length aligned to only bacterial, virus, chloroplast or mitochondrion sequences were removed. Additionally, contigs shorter than 50 kb and having a read coverage lower than 20× or higher than 200× were also excluded in the final assembly. The contigs were then clustered into 90 haplotype groups (six haplotypes per each of the 15 linkage groups, corresponding to 15 chromosomes) based on simplex markers from a phased genetic map constructed with a full-sib hexaploid sweetpotato population derived from a cross between the cultivars ‘Beauregard’ and ‘Tanzania’ (Mollinari et al., 2020). The flanking sequences of simplex SNP markers were aligned to the contigs using bowtie (v1.1.2) (Langmead et al., 2009), requiring proper-paired end-to-end alignment. Genotypes at the simplex markers on a contig were used to assign the contig to a specific haplotype. Chimeric contigs were identified when haplotype switches were indicated by the simplex markers and the switch regions coincided with a depletion of Hi-C contact signals. The breakpoints on chimeric contigs were determined with the misjoin detection strategy within the 3D-DNA suite (Dudchenko et al., 2017).

The construction of the 90 pseudochromosomes involved two main steps: (1) grouping the fully phased contigs into 90 haplotype groups and (2) anchoring of the contigs to chromosomes within each haplotype. The hifiasm primary contigs were used to build a 15-chromosome consensus monoploid reference assembly, disregarding phase information. These primary contigs were aligned against each other using minimap2 (v2.21-r1071) (Li, 2021), and the longest non-allelic contigs were anchored to the 15 chromosomes using the hexaploid sweetpotato genetic map (Mollinari et al., 2020). This consensus assembly was then used to sort the phased contigs into 15 chromosomes and provide synteny information for chromosome construction. The phased contigs were aligned to the consensus assembly with minimap2 (v2.21-r1071) and assigned to the 15 chromosomes. Approximately 90% of the fully phased assembly was grouped into 90 haplotypes based solely on genetic information. Hi-C contact information was then used to assign the remaining phased contigs to the appropriate haplotypes. Paired-end Hi-C reads were aligned to the phased contigs using BWA-MEM (v0.7.17) (Li, 2013) with the parameter ‘-5SP’. Uniquely mapped reads with a mapping quality ≥ 20 were retained. Contact links between contigs derived from read pairs were extracted from the Hi-C read alignments using hickit (https://github.com/lh3/hickit). Examining the Hi-C contact links among haplotype-resolved contigs via simplex markers showed that a contig had the highest Hi-C contact signals with other contigs belonging to the same haplotype (**Supplementary Fig. 20**). Therefore, a contig’s average Hi-C contact frequencies with the 90 haplotype groups, combined with synteny and allelic information, were used to assign the contig to a specific haplotype (**Supplementary Fig. 21**). The phased genetic maps, Hi-C data, and synteny with the consensus assembly were used to construct the pseudochromosomes. Hi-C reads were aligned to the phased contigs using Juicer (Durand et al., 2016), and the resulting duplicate-free list of paired alignments was used as input for 3D-DNA (Dudchenko et al., 2017) scaffolding within each haplotype. Scaffolding of contigs based on their synteny with the consensus assembly was performed using RagTag (Alonge et al., 2022). The ordering and orientation of the contigs based on these three lines of evidence, genetic maps, Hi-C scaffolding result, and synteny, were combined using ALLMAPS (Tang et al., 2015).

### Assembly quality evaluation

Merqury (v1.3) (Rhie et al., 2020) was used to evaluate the base accuracy and completeness of the assembly. The completeness was also evaluated by comparing the k-mer spectrum in HiFi reads and the phased assembly using the K-mer Analysis Toolkit (KAT) (v2.4.2) (Mapleson et al., 2017). The completeness in terms of capturing conserved plant genes was assessed with BUSCO (v5.4.4) (Simão et al., 2015) using 1,614 genes from the ‘embryophyta_ odb10’ database. Assembly contiguity was assessed using LTR Assembly Index (LAI) (Ou et al., 2018). To assess the correctness of haplotype phasing at the chromosome level, we aligned Hi-C reads to the assembly and the resulting contact maps were visualized with HiCExplorer (v3.7.2) (Wolff et al., 2018). To further assess haplotype phasing, we constructed a linkage map using the Beauregard × Tanzania population described in Mollinari et al. (2020) with the phased ‘Tanzania’ genome assembly as the reference. Variant and genotype calling were performed using GATK (Mckenna et al., 2010), treating each phased haplotype separately. The linkage map was constructed using MAPpoly (Mollinari and Garcia, 2019), resulting in a phased ‘Tanzania’ genetic map comprising 41,483 SNPs with a total length of 1,623 cM. The density of reference alleles of SNPs called within each haplotype across 2-Mb non-overlapping windows throughout the ‘Tanzania’ genome was used to evaluate the correctness of haplotype phasing (**Supplementary Fig. 8**).

### Genome annotation

Repeats were first identified in the ‘Tanzania’ genome assembly with RepeatModeler (v2.03) (Flynn et al., 2020) and protein-coding genes were filtered out from the identified repeats using ProtExcluder (v1.2) (Campbell et al., 2014) to create a custom repeat library (CRL). This CRL was then combined with Viridiplantae repeats from RepBase (v20150807) (Bao et al., 2015) to generate the final CRL. Repeat masking of the ‘Tanzania’ genome assembly was performed using the final CRL and RepeatMasker (v4.1.2-p1) (Tarailo-Graovac & Chen, 2009) with the parameters ‘-e ncbi -s -nolow -no_is -gff’.

‘Tanzania’ RNA-seq reads from Wu et al. (2018) were processed using Cutadapt (v4.6) (Martin 2011) with a minimum length of 70 nt and a quality cutoff of 10, and then aligned to the ‘Tanzania’ genome assembly using HISAT2 (v2.2.1) (Kim et al., 2019). Additionally, full-length cDNA libraries were prepared from immature leaf, storage roots, and fibrous roots of ‘Tanzania’ using the Oxford Nanopore Technologies (ONT) PCR-cDNA Barcoding kit (SQK-PCB109) and sequenced on FLO-MIN106 Oxford Nanopore (ONT) flow cells. ONT cDNA reads were processed with Pychopper (v2.5.0; https://github.com/nanoporetech/pychopper) and trimmed reads longer than 500 nt were aligned to the ‘Tanzania’ genome using minimap2 (v2.17-r941) (Li 2021) with a maximum intron length of 5,000 nt. The aligned RNA-seq and ONT cDNA reads were each assembled using Stringtie (v2.2.1) (Kovaka et al., 2019), and assembled transcripts shorter than 500 nt were removed.

Initial gene models were generated using BRAKER2 (v2.1.6) (Brůna et al., 2021) using the soft-masked genome assembly and RNA-seq read alignments as hints. These gene models were then refined through two rounds of PASA2 (v2.5.2) (Haas et al., 2008) to create a working gene model set. High-confidence gene models were identified from the working gene model set by filtering out those without expression evidence, a Pfam domain match, or those that were partial gene models or contained an interior stop codon. Functional annotation of the predicted genes was performed by searching their protein sequences against the TAIR (v10) (Lamesch et al., 2012) and Swiss-Prot plant protein (release 2015_08) databases using BLASTP (v2.12.0) (Altschul et al., 1990) and the Pfam (v35.0) (El-Gebali et al., 2019) database using PfamScan (v1.6) (Li et al., 2015). Functional enrichment of genes was performed using Blast2GO (Götz et al., 2008).

### Variant calling

Using haplotype A of ‘Tanzania’ as the reference, the other five haplotypes were each aligned to the reference haplotype for variant calling. Alignment between haplotypes was performed using the AnchorWave software (Song et al., 2022). SNPs were called using BCFtools (Li, 2011). SVs were identified in the aligned genomic regions with NucDiff (Khelik et al., 2017). SVs were then merged, and redundant ones were removed with findDup.R according to instructions provided on GitHub (https://github.com/vgteam/giraffe-sv-paper/blob/master/scripts/sv).

### Construction of syntenic orthologous gene groups

Orthologous and paralogous gene groups were determined among *I. trifida* NCNSP0306 and the six haplotypes of ‘Tanzania’ using OrthoFinder (v2.5.5) (Emms and Kelly, 2019). High-confidence genomic syntenic blocks between *I. trifida* and each ‘Tanzania’ haplotype, as well as between pairs of haplotypes were identified using MCScanX (Wang et al., 2012) with an E value cutoff of 1e-10, which was used to classify the OrthoFinder-defined orthologous and paralogous gene groups into 1) syntenic gene groups, 2) orthologous groups without synteny information, and 3) singletons. *K*_S_ values of homologous gene pairs were calculated using the Yang-Nielsen algorithm implemented in the PAML package (Yang et al., 1997). Synteny map was generated with SynVisio (https://synvisio.github.io/#/) based on the MCScanX results.

### Ancestry inference

Illumina whole-genome resequencing data of two *I. aequatoriensis* accessions (CH81.2 and CH71.3) and one *I. trifida* accession (CIP460663) from Muñoz-Rodríguez et al. (2022), eight *I. aequatoriensis* (PI561246, PI561247, PI561248, PI561255, PI561258, PI561261, K300_1, K300_5) and eight *I. batatas* 4× accessions (ECAL_2145(1)_1, ECAL_2156(1)_1, ECAL_2232(3)_2, ECAL_2156(10)_2, ECAL_2156(5)_1, ECAL_2192_2, ECAL_2262_1, ECAL_2293(2)_1) from Yan et al. (2024), and one *I. trifida* accession (NCNSP0306) from Wu et al. (2018) (**Supplementary Table 12**), were used for ancestry inference. Raw sequencing reads were processed with Trimmomatic (v0.36) (Bolger et al., 2014) to remove adaptors and low-quality sequences. Ancestry inference of ‘Tanzania’ was based on both genetic distance and sequence identity between sweetpotato and the wild relatives (**Supplementary Fig. 22**). To calculate the genetic distance between each tetraploid wild accession and ‘Tanzania’, SNPs were called based on read alignments. Briefly, cleaned Illumina reads were mapped to each haplotype in the ‘Tanzania’ genome using BWA-MEM (v0.7.17) (Li, 2013) with default parameters, followed by duplicate alignment marking using SAMBLASTER (v0.1.26) (Faust and Hall, 2014). The processed alignment bam files were then used for SNP calling with the Sentieon software package (https://www.sentieon.com/), which was built based on GATK (Mckenna et al., 2010). Variants from each sample were called using the Haplotyper function in Sentieon with ‘--ploidy 4’, followed by joint variant calling with the GVCFtyper function. Raw SNPs were filtered using GATK (v3.8) with parameters ‘QD<2.0 || FS>60.0 || MQ<40.0 || SOR>3.0 || MQRankSum<−12.5 || ReadPosRankSum<−8.0’. A phylogenetic tree of six sweetpotato haplotypes, ten wild *I. aequatoriensis* and eight *I. batatas* 4× tetraploid accessions was constructed for each ‘Tanzania’ genomic window containing 5,000 non-overlapping SNPs using IQ-TREE (v.2.3.1) (Nguyen et al., 2015) with the nucleotide substitution model TVMe+ACS+R2 and 100 bootstrap replicates. The resulting genetic distance matrices were used to classify the sweetpotato genomic windows as either *I. aequatoriensis*-like or *I. batatas* 4×-like using the k-nearest neighbors (KNN) algorithm with seven nearest wild accession neighbors of each sweetpotato haplotype. Ancestry was determined when at least four out of seven nearest neighbors were from the same wild species. To analyze sequence similarity between a wild relative and ‘Tanzania’, Illumina read mapping and variant calling followed the same procedures as above, except that the phased ‘Tanzania’ genome was used as the reference and variants were called using the Sentieon Haplotyper function with ‘- -ploidy 1’. For each ‘Tanzania’ genomic window, sequence identity between a wild accession and ‘Tanzania’ was calculated as the number of sites with the same genotype as ‘Tanzania’ divided by the total number of genotyped sites in the window. Windows with fewer than 10,000 genotyped sites were excluded from the analysis. Introgressed sequences in ‘Tanzania’ were identified as those with high sequence identities (>99.9%) to *I. batatas* 4× individuals, corresponding to the small *I. batatas* 4×-specific peak near 100% in the identity distribution plot, distinct from the main *I. batatas* 4×-like identity distribution peaking at 99.3% (**Fig. 2a**).

## Data availability

Raw genome sequencing reads have been deposited in the NCBI BioProject database under the accession no. PRJNA1138727. The ‘Tanzania’ phased and consensus genome assemblies, and annotated genes are also available at Sweetpotato Genomics Resource (http://sweetpotato.uga.edu/).

## Acknowledgements

We thank Dr. Sharon Williamson from NC State University for her assistance in sample collection and Dr. Benard Yada from National Crops Resources Research Institute, Uganda for providing the ‘Tanzania’ images. This research was supported by grants from the Bill and Melinda Gates Foundation through SweetGAINS (OPP1213329) and RTB Breeding (CGIAR Investment ID1523-BMGF) Projects under a subcontract with the International Potato Center, Lima, Peru, and USDA National Institute of Food and Agriculture (2022-67013-36269).

## Author contributions

Z.F., C.R.B. and G.C.Y. designed and managed the project. S.W. and Z.F. coordinated the genome sequencing. S.W. and H.S. performed the genome assembly and evaluation. J.P.H. and M.K. performed ONT transcriptome sequencing and genome annotation. S.W., H.S., M.Y., H.W. and J.Y. analyzed the origin of sweetpotato. M.M. and G.D.S.G. constructed the phased genetic map. S.W. wrote the manuscript. Z.F. revised the manuscript. All authors read and approved the manuscript.

## Competing interests

The authors declare no competing interests.

## Notes

### Competing Interest Statement

The authors have declared no competing interest.

