## Supplementary Figures for "Phased chromosome-level genome assembly provides insight into the origin of hexaploid sweetpotato"

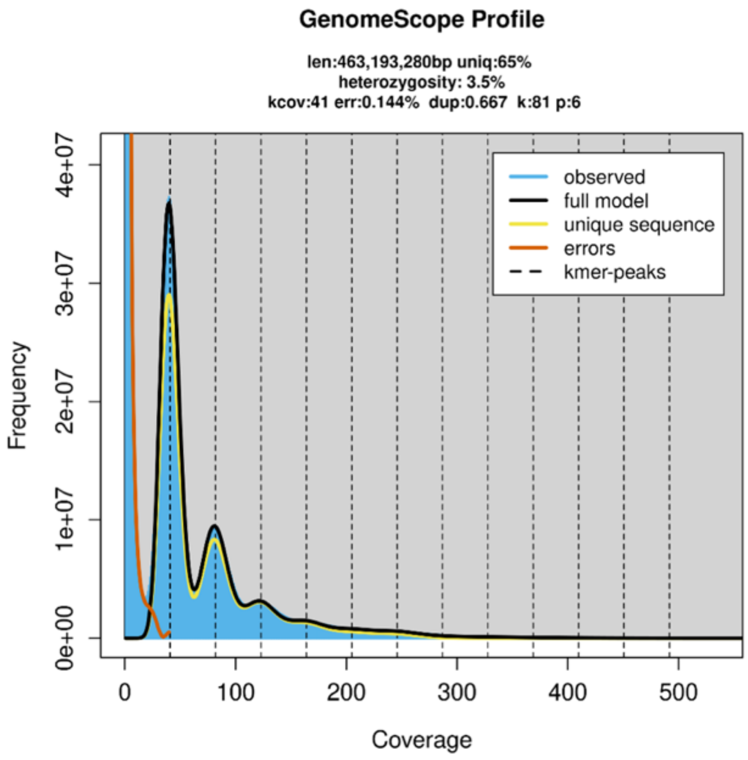


**Supplementary Figure 1** *I. batatas* 'Tanzania' genome size estimation using k-mer analysis. K-mer abundance in the HiFi reads was calculated using JellyFish (https://github.com/gmarcais/Jellyfish) with a k-mer size of 81. Genome size was estimated using GenomeScope 2.0 (https://github.com/tbenavi1/genomescope2.0). The estimated monoploid genome size is 463.2 Mb; thus, the estimated hexaploid genome size of 'Tanzania' is 2779.2 Mb.

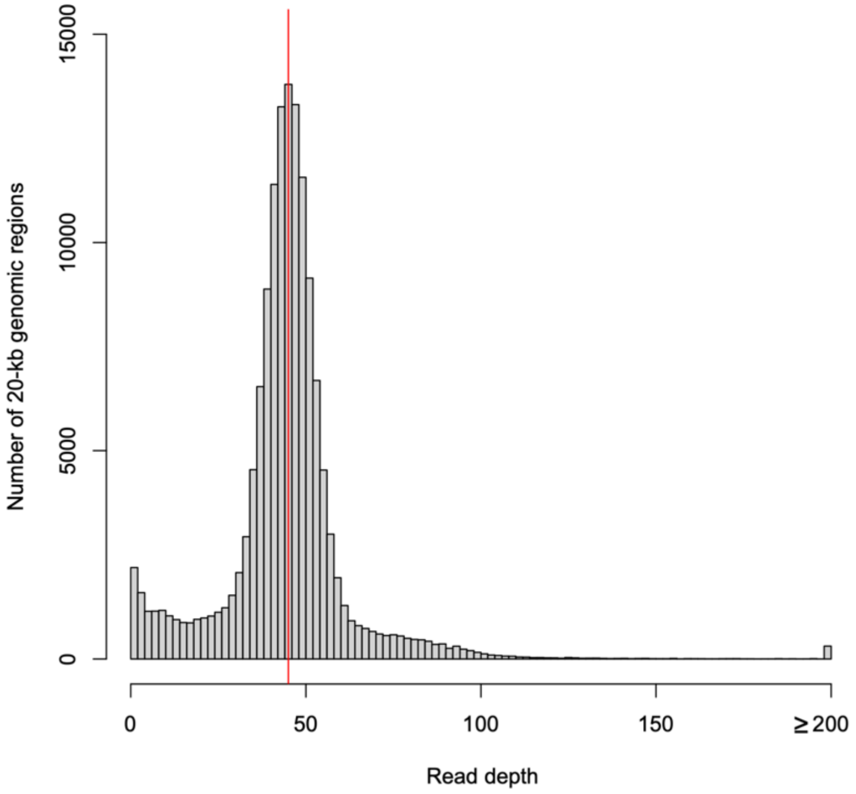


**Supplementary Figure 2** Distribution pattern of HiFi read coverage depth on the phased 'Tanzania' genome assembly.

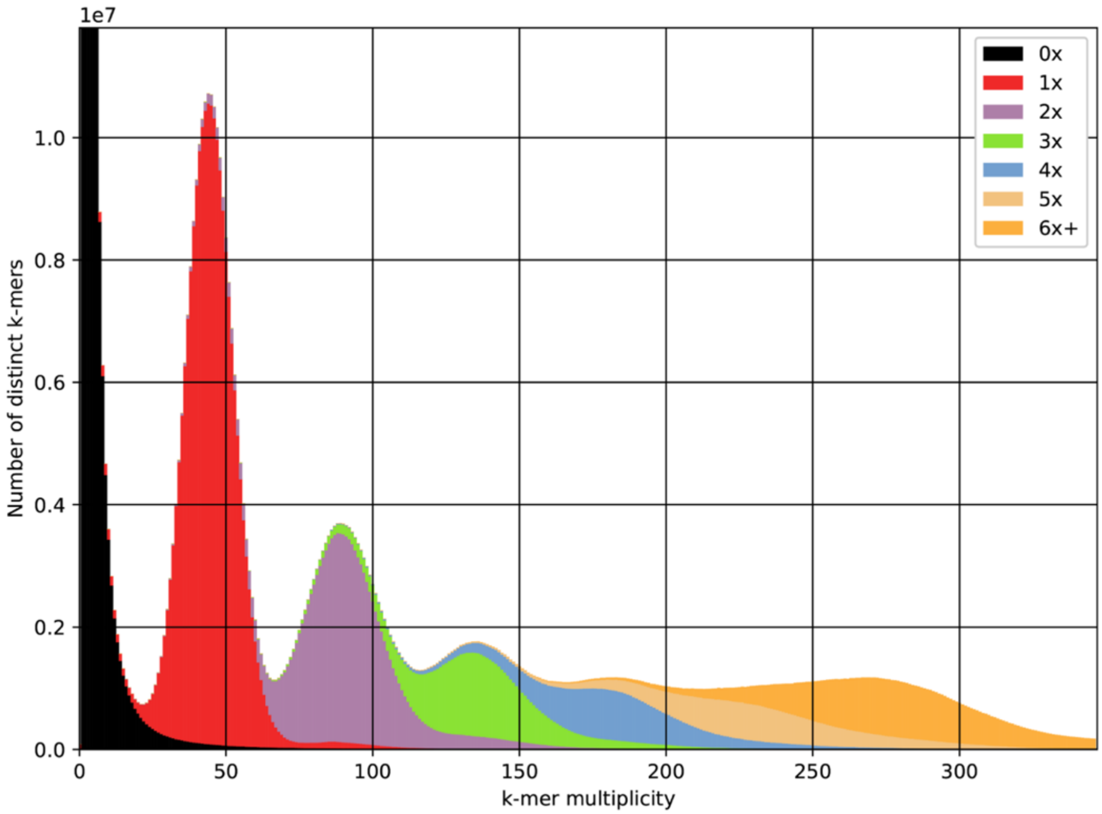


**Supplementary Figure 3** Comparison of 21-mer spectra between HiFi reads and the phased 'Tanzania' assembly. Black indicates 21-mers present in HiFi reads but absent from the assembly. Red, purple, green, blue, yellow and orange denote 21-mers occurring once, twice, three, four, five and six times, respectively, in the assembly.

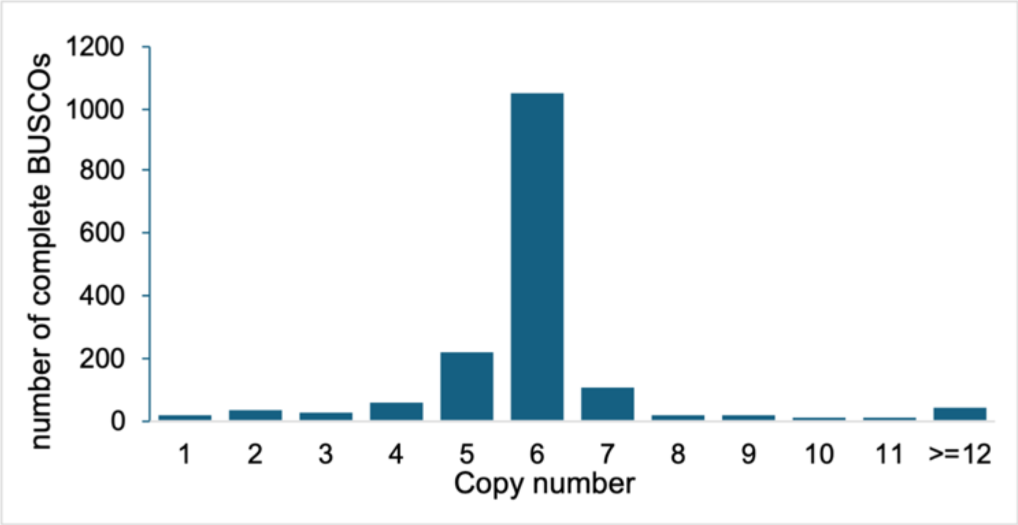


**Supplementary Figure 4** Distribution of copy numbers of complete BUSCO genes captured in the phased ‘Tanzania’ genome assembly.

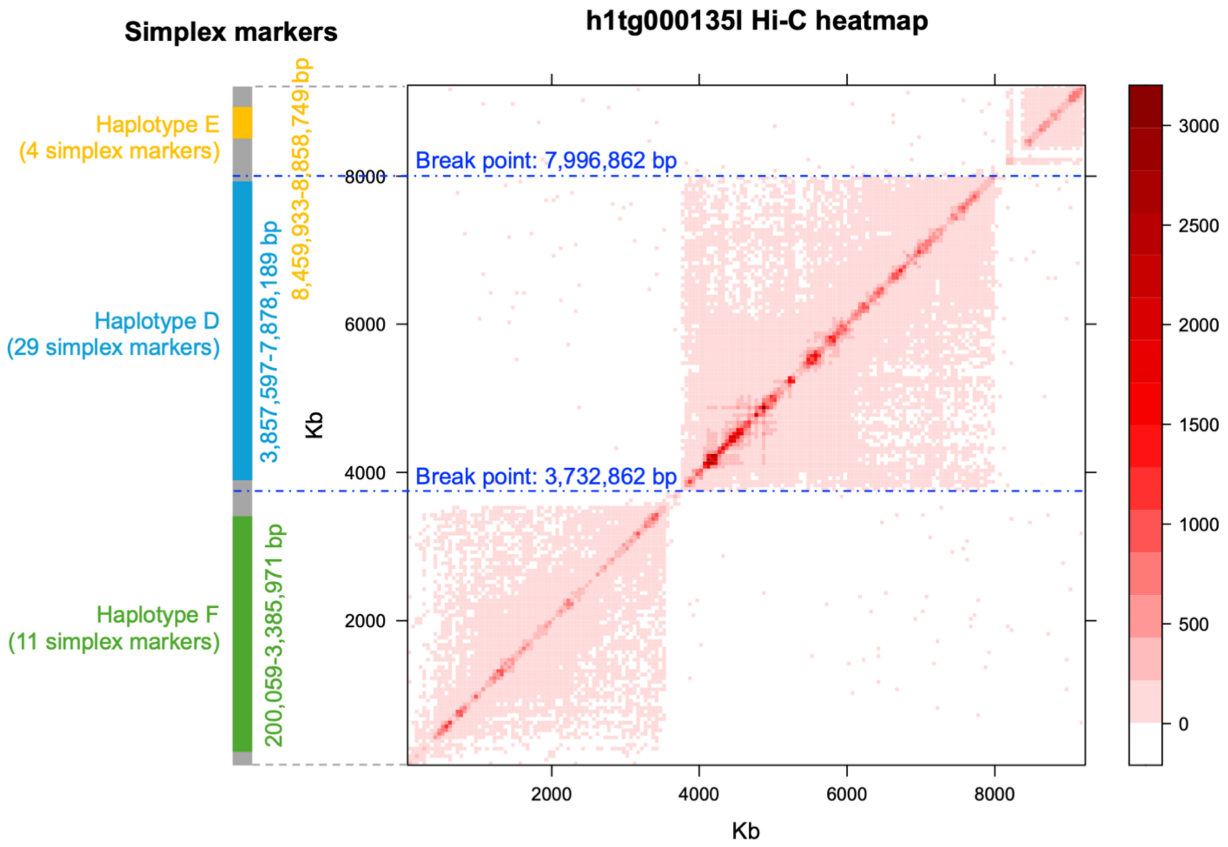


**Supplementary Figure 5** Detection and correction of a chimeric contig, h1tg000135l, based on genetic markers and Hi-C contact signals. The bar on the left shows the positions and numbers of simplex markers supporting three different haplotypes across three regions in the same contig. Gray bars indicate regions not covered by simplex markers. Phase switching sites indicated by the genetic markers coincide with regions with depleted Hi-C contact signals, as shown in the heatmap on the right. Exact break points were determined using the misjoin detection algorithm implemented by 3D-DNA (https://github.com/aidenlab/3d-dna).

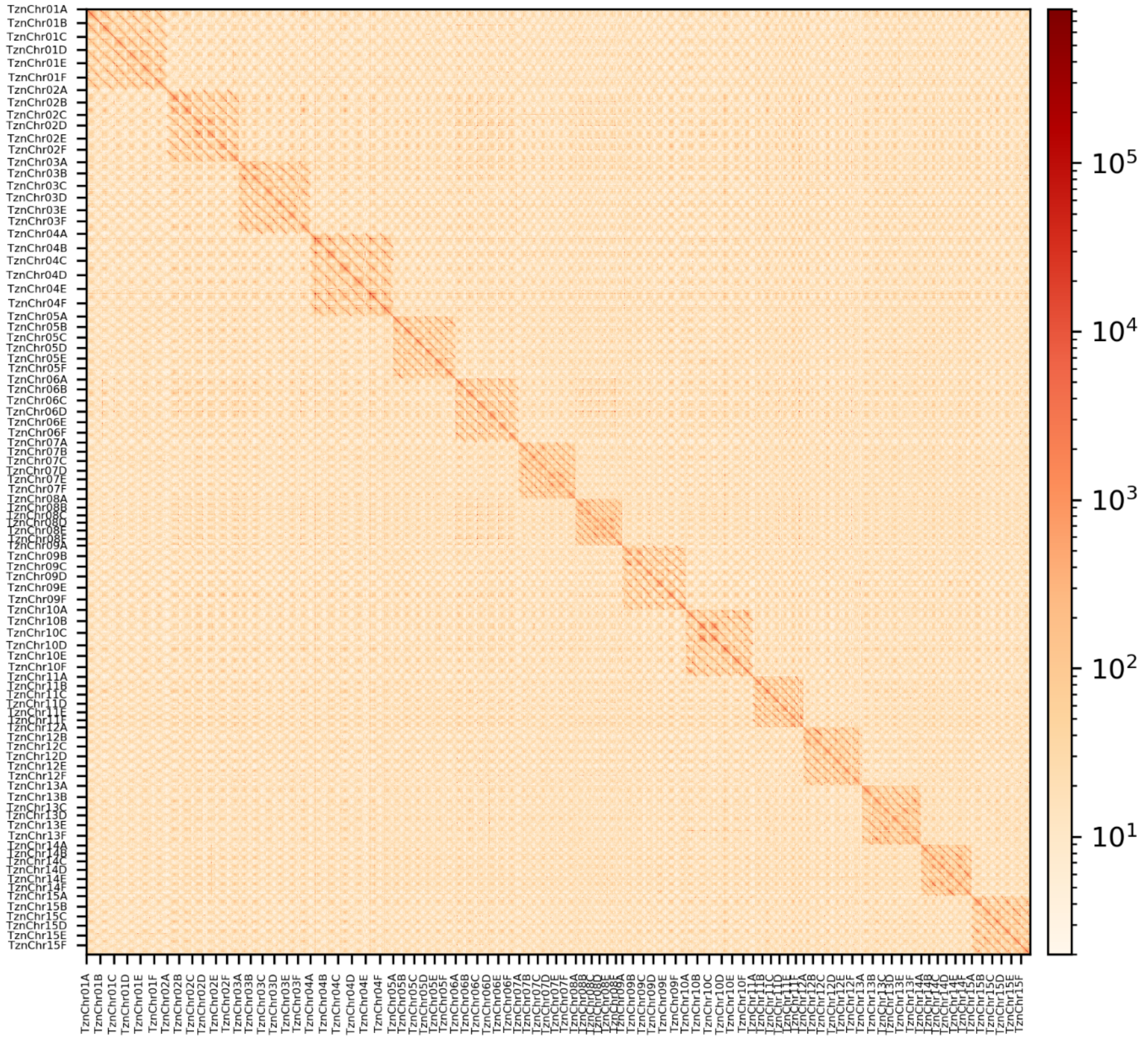


**Supplementary Figure 6** Hi-C contact heatmaps for the 90 chromosomes in the 'Tanzania' assembly. The Hi-C contact heatmap was generated using all mapped reads. Each of the 15 homoeologous chromosome groups forms a distinct block.

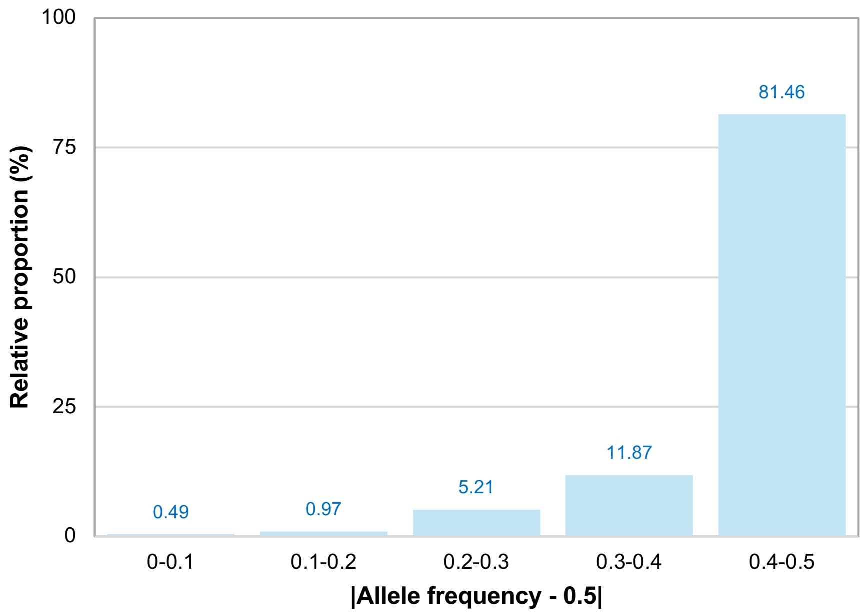


**Supplementary Figure 7** Validating the accuracy of genome phasing. Allele frequency was used to verify the accuracy of the phasing of the 'Tanzania' genome assembly using nPhase (https://github.com/OmarOakheart/nPhase). An allele frequency close to 0.5 or the absolute value of the allele frequency minus 0.5 (|Allele frequency – 0.5|) close to zero is considered a potential incorrectly phased region. A very small portion (0.49%) of the assembled 'Tanzania' genome had |Allele frequency – 0.5| close to zero, indicating a very low potential phase switching error rate in the 'Tanzania' genome assembly.

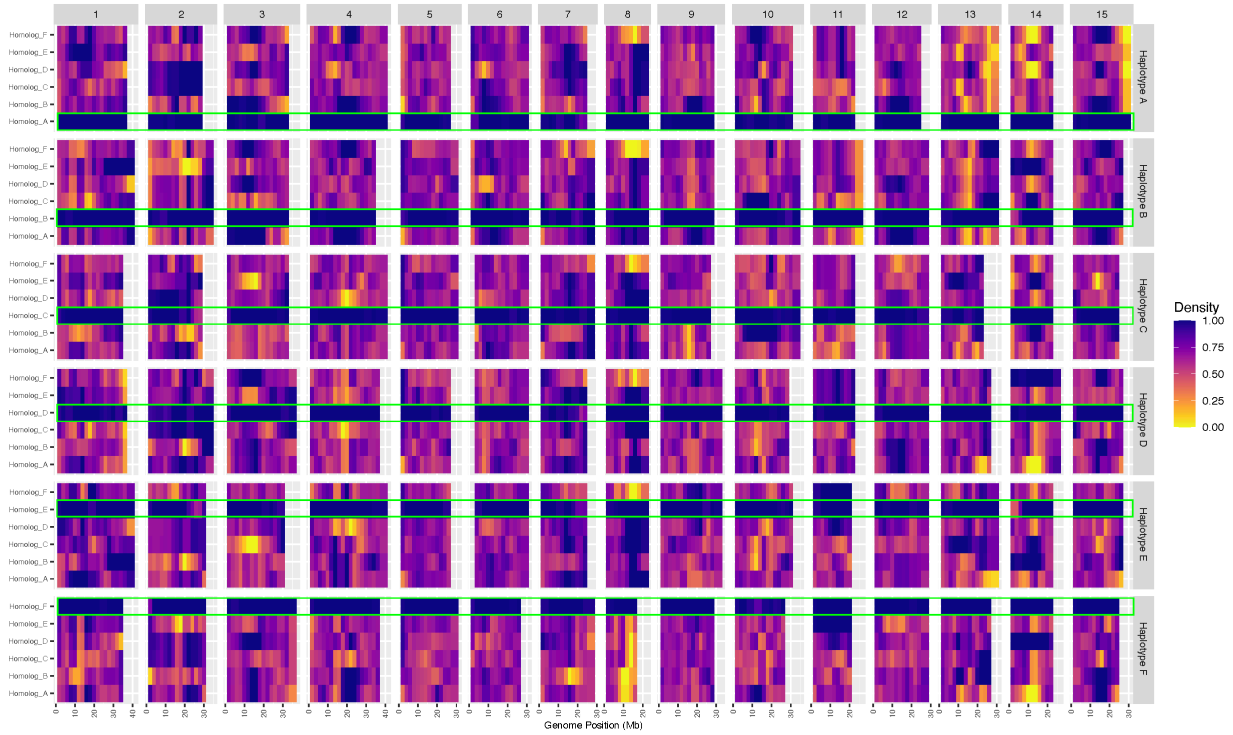


**Supplementary Figure 8** Haplotype phasing assessment of the 'Tanzania' assembly using the phased genetic map. The density of reference alleles of SNPs within each haplotype across 2-Mb non-overlapping windows throughout the 'Tanzania' genome assembly is shown. For each chromosome and haplotype combination used to obtain the SNPs, the majority of reference alleles in these SNPs (98.6%) were phased within a specific haplotype, as indicated by the green boxes.

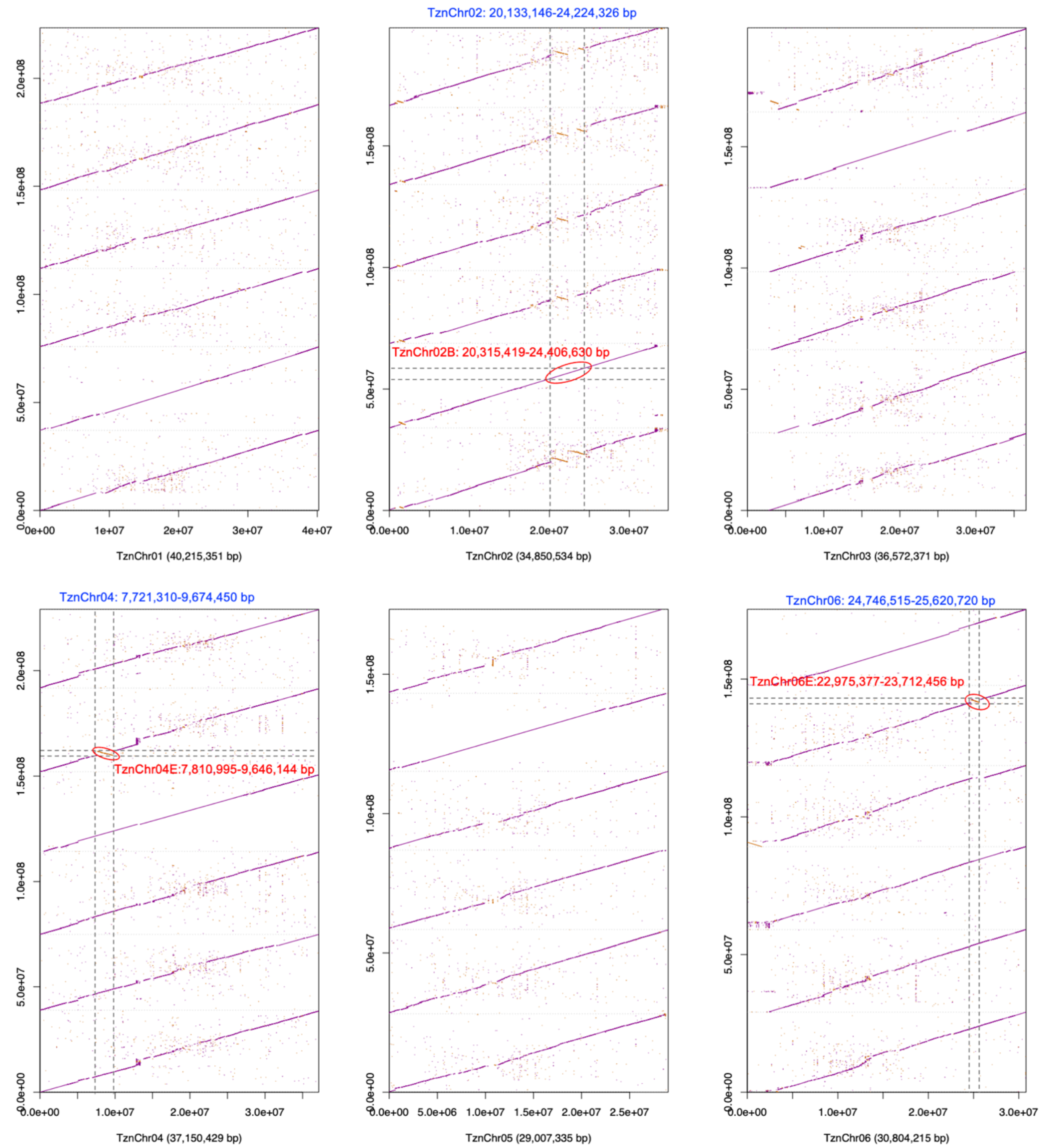


**Supplementary Figure 9** Synteny between the six phased 'Tanzania' haplotypes and the consensus chromosomes. Large inversions are indicated with red circles. Genomic positions of the inversions in the phased and consensus assemblies are shown in red and blue, respectively.

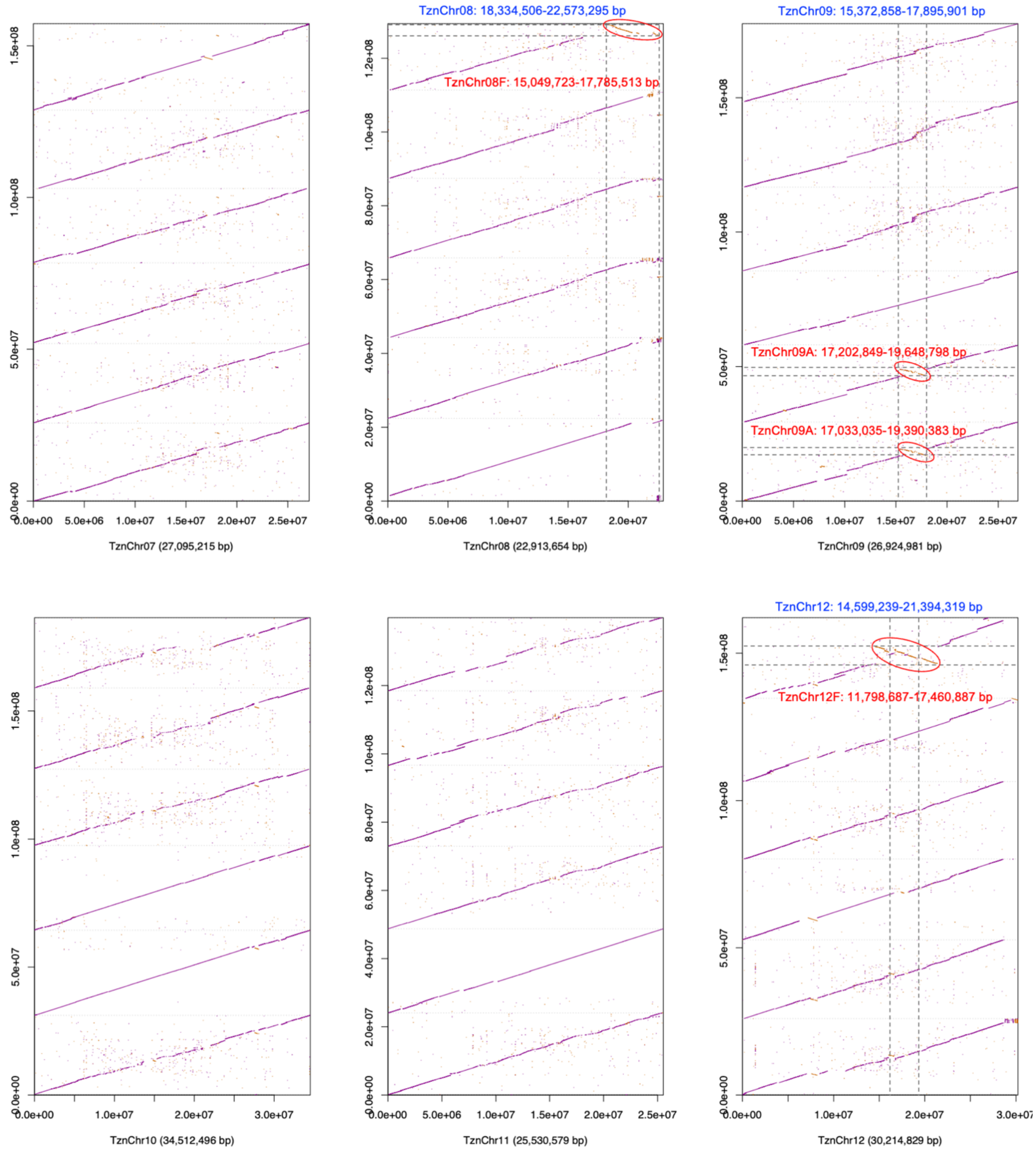


**Supplementary Figure 9** Continued

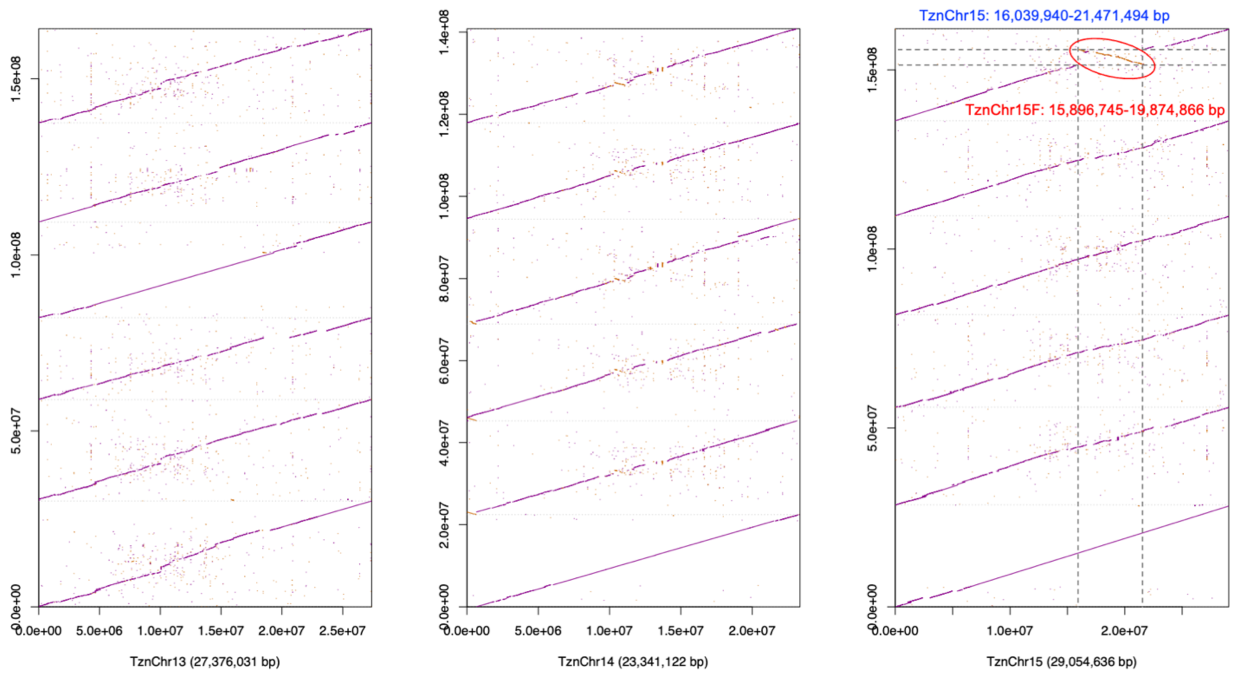


**Supplementary Figure 9** Continued

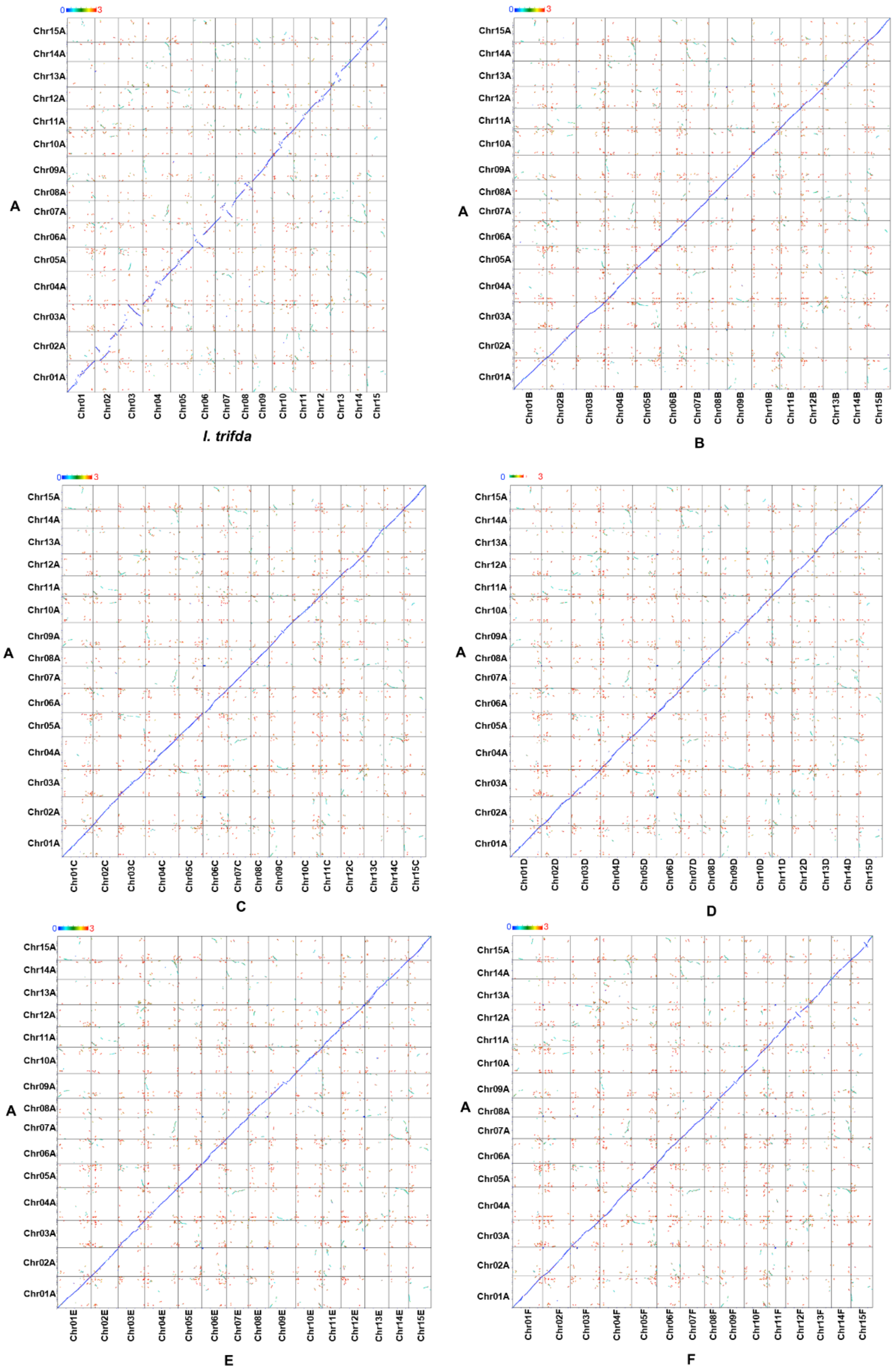


**Supplementary Figure 10** Synteny between ‘Tanzania’ haplotype A and *I. trifida* NCNSCP0306 genome, as well as between ‘Tanzania’ haplotype A and other five haplotypes. Syntenic gene blocks are labeled with colors representing the medium $K_S$ values.

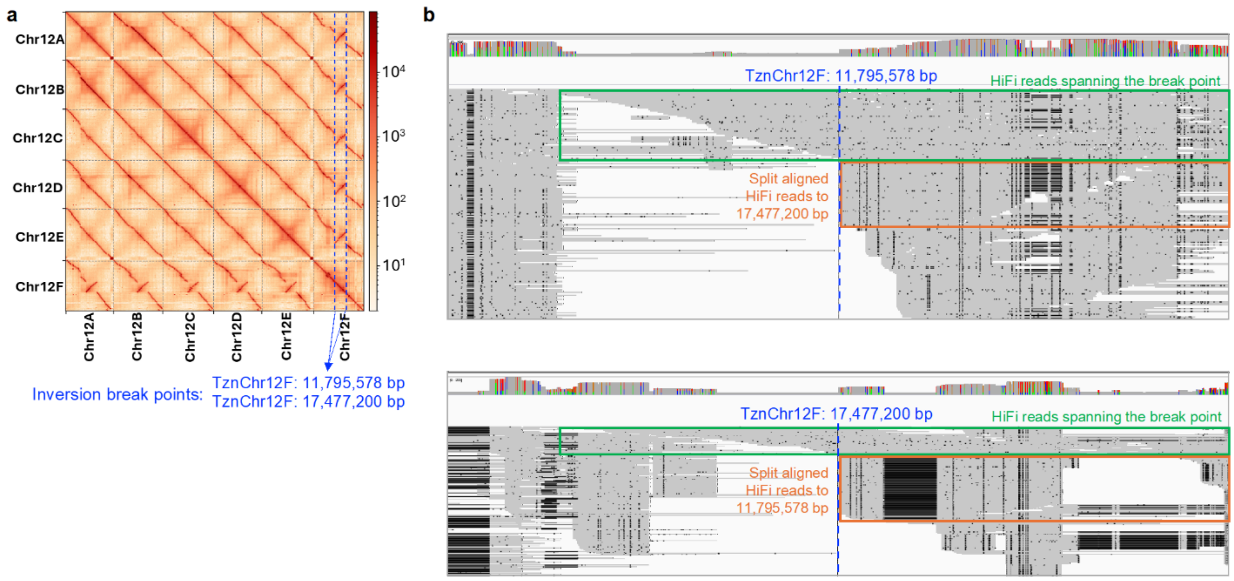


**Supplementary Figure 11** Large inversions among the six 'Tanzania' haplotypes supported by Hi-C contact signals and HiFi read alignments. The example shown here is the 5.68-Mb inversion on 'Tanzania' chromosome 12F. **a**, Heatmap of the Hi-C contact signals among the six haplotypes of 'Tanzania', supporting the inversion on chromosome 12F. **b**, HiFi read alignments using TznChr12F as the reference. Reads spanning the inversion breakpoints at TznChr12F 11.8 Mb and 17.5 Mb, which originated from haplotype TznChr12F, are highlighted by the green boxes. Reads that are split-aligned, which originated from the other five haplotypes, are highlighted by orange boxes.

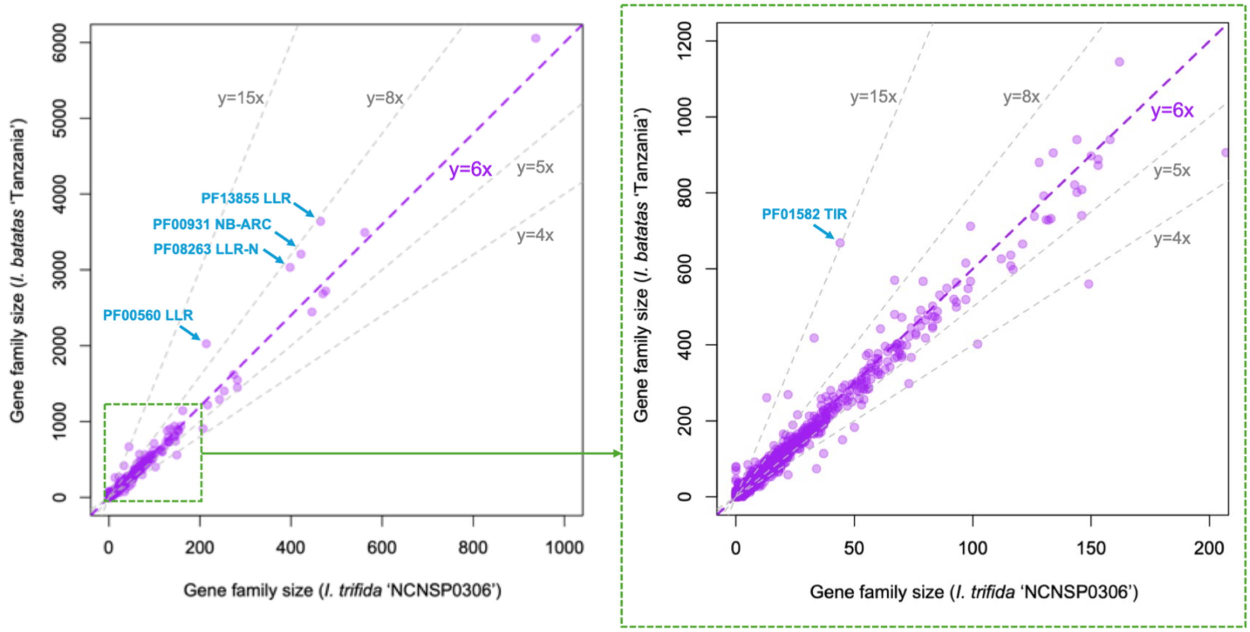


**Supplementary Figure 12** Dotplot showing the fold change of gene family sizes between the *I. trifida* NCNSP0306 and phased *I. batatas* ‘Tanzania’ genome assemblies. An enlarged view of gene families with sizes smaller than 200 is shown on the right.

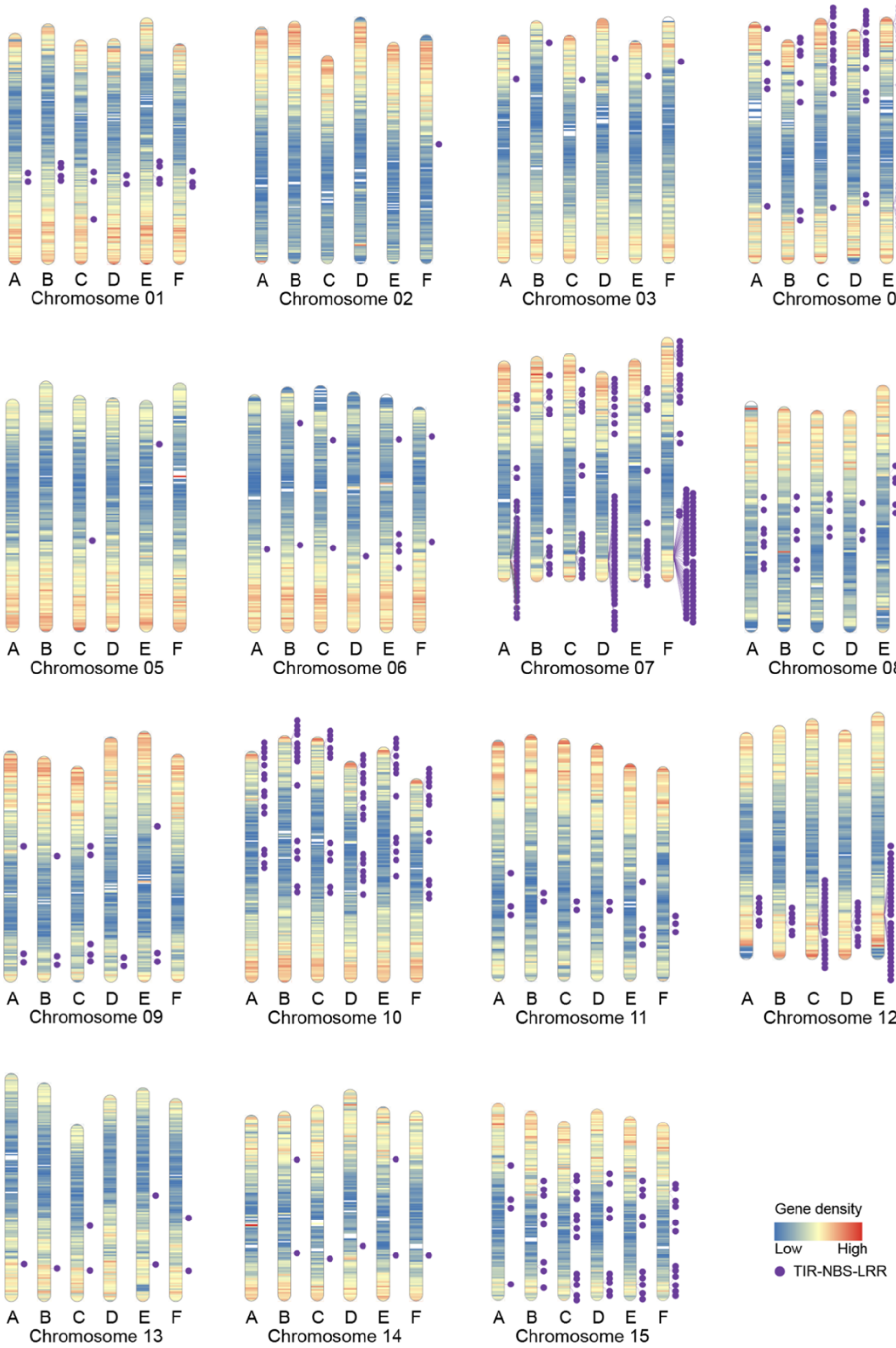


**Supplementary Figure 13** Distribution of TIR-NBS-LRR genes across the phased 'Tanzania' genome.

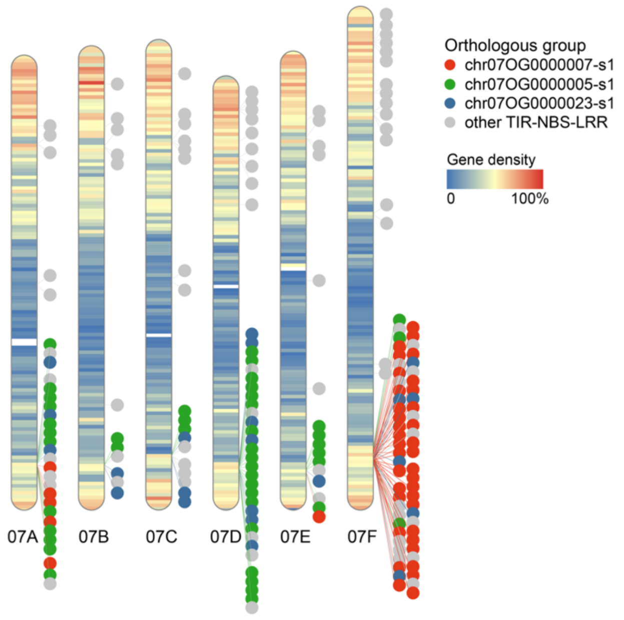


**Supplementary Figure 14** Distribution of TIR-NBS-LRR genes across the six haplotypes of 'Tanzania' chromosome 7. Genes within the same syntenic paralogous group are labeled with the same color.

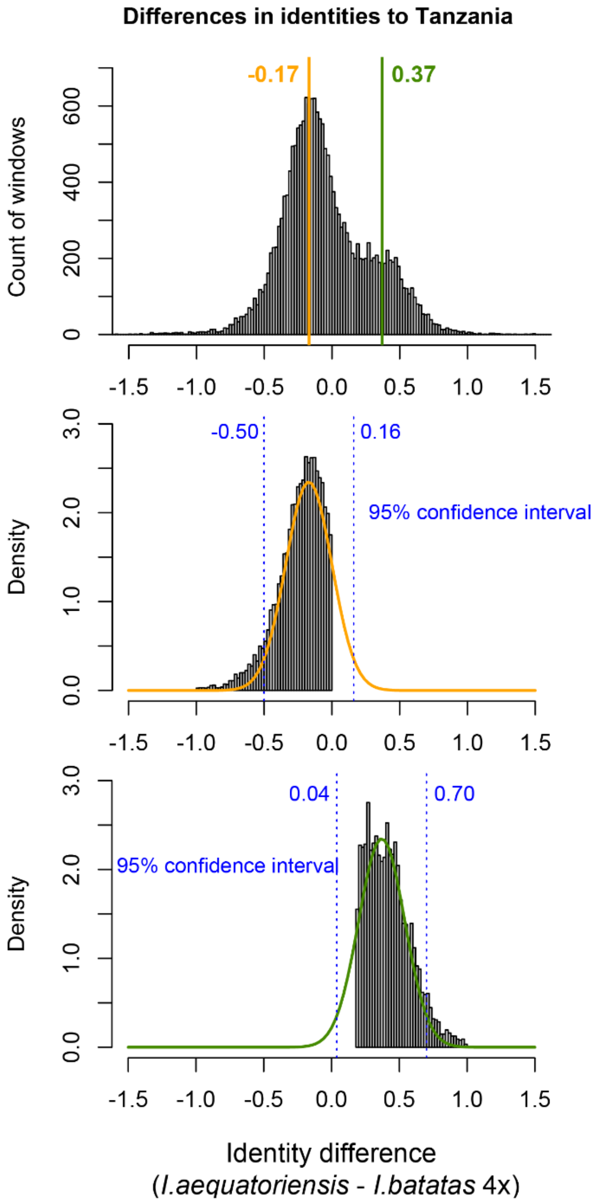


**Supplementary Figure 15** Distribution of sequence identity differences between the two wild tetraploid species and hexaploid sweetpotato. For each analyzed phased genomic window, the sequence identity difference was calculated as: the mean sequence identity between ten *I. aequatoriensis* accessions and the hexaploid sweetpotato ‘Tanzania’ minus the mean sequence identity between eight *I. batatas* 4× accessions and the hexaploid sweetpotato ‘Tanzania’. The two peaks of the identity difference distribution are labeled by the orange and green lines in the top panel. Normal distributions were fitted to the patterns displayed by ‘Tanzania’ genomic windows with higher sequence identity to *I. batatas* 4× (middle) and those with higher identity to *I. aequatoriensis* (bottom). The 95% confidence intervals are indicated by the blue dotted lines.

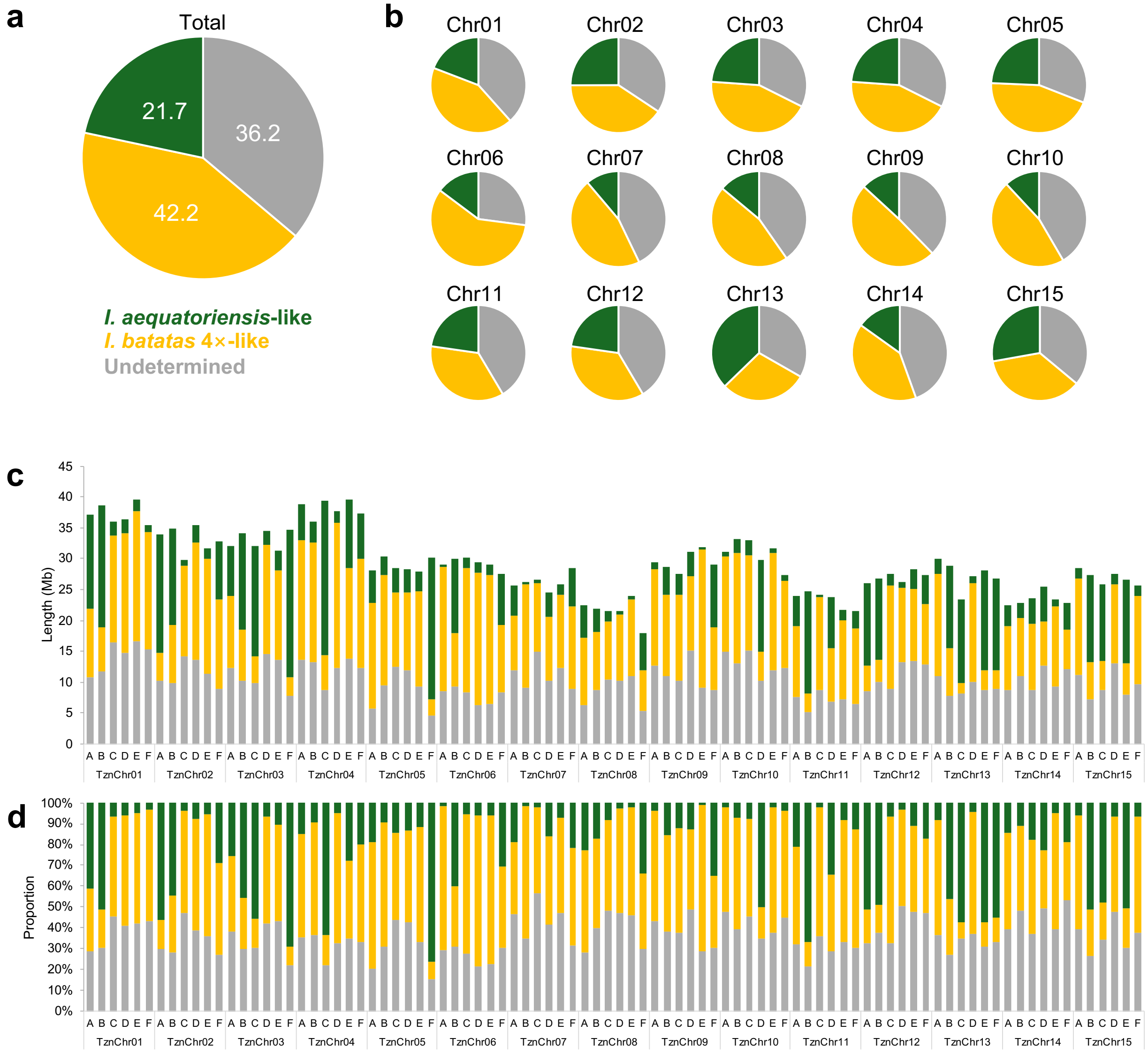


**Supplementary Figure 16** Proportion and length of 'Tanzania' genomic sequences inferred to have different ancestry types. **a**, Proportion calculated across the entire genome. **b**, Proportions calculated for each homoeologous chromosome group. **c**, Lengths of 'Tanzania' sequences with different ancestry types across the 90 chromosomes. **d**, Proportion of 'Tanzania' sequences with different ancestry types in each of the 90 chromosomes.

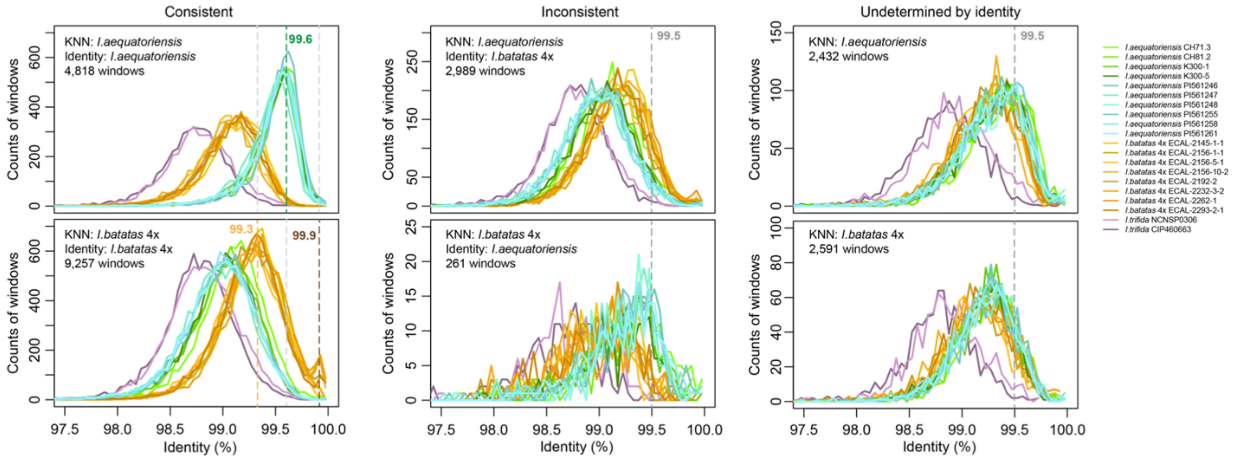


**Supplementary Figure 17** Distribution of sequence identities between 'Tanzania' and wild *I. aequatoriensis*, *I. batatas* 4×, and *I. trifida* accessions in 'Tanzania' genomic windows with inferred ancestry based on genetic distance and sequence identity. Windows with consistent and contradictory ancestry inference between the two methods are shown on the left and in the middle, respectively. Windows whose ancestry could not be determined based on sequence identity are shown on the right. KNN refers to ancestry inference based on genetic distance using a k-nearest neighbor algorithm.

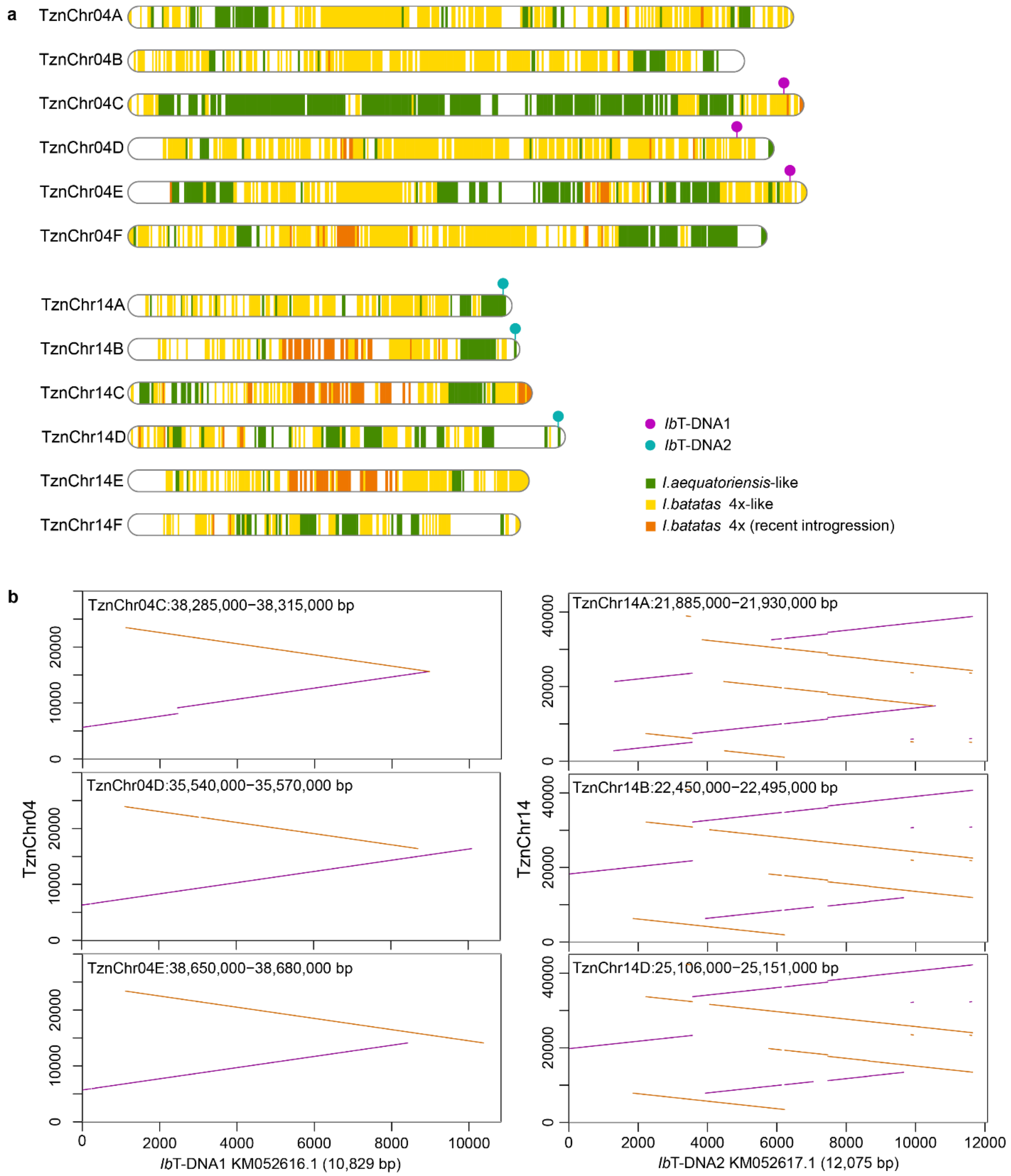


**Supplementary Figure 18** *Ib*T-DNA sequences in the hexaploid sweetpotato 'Tanzania' genome. **a**, Positions of *Ib*T-DNA1 and *Ib*T-DNA2 sequences on the 'Tanzania' chromosomes. **b**, Alignments between the *Ib*T-DNAs and the 'Tanzania' genomic regions containing these *Ib*T-DNA insertions.

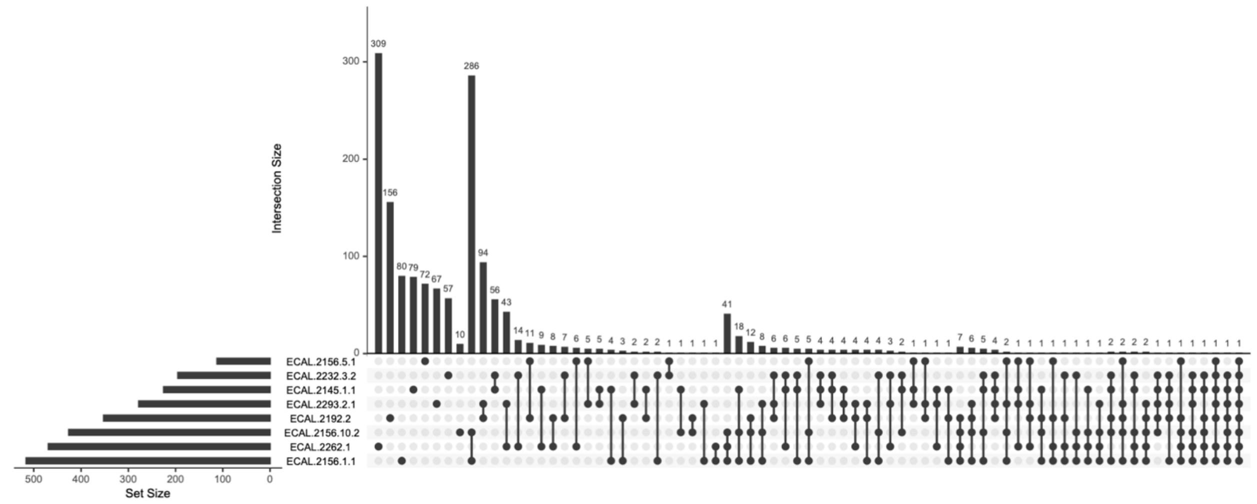


**Supplementary Figure 19** UpSet plot showing the number of genomic regions introgressed from wild *I. batatas* 4× accessions into the hexaploid sweetpotato.

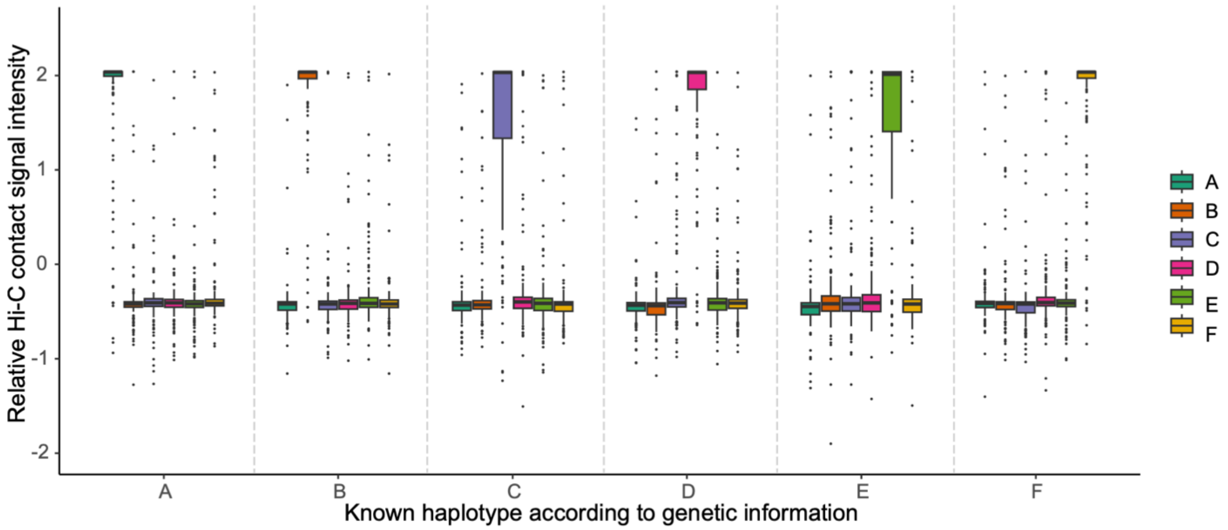


**Supplementary Figure 20** Relative Hi-C contact signal intensities of contigs compared to those within the same haplotype and those in other haplotypes.

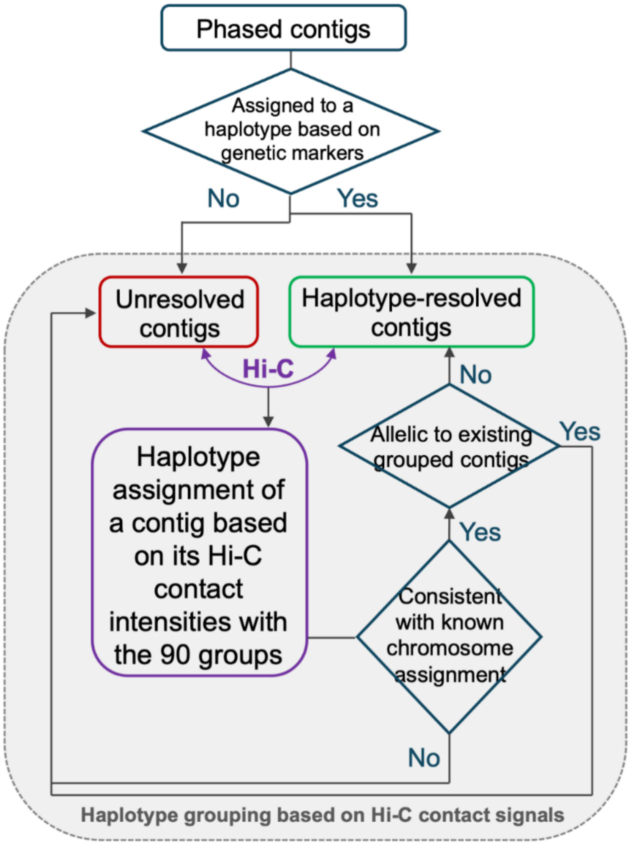


**Supplementary Figure 21** Workflow for haplotype phasing of contigs using genetic and Hi-C contact information. First, genetic information was used to assign the phased contigs to 90 groups. The remaining contigs were then assigned to these 90 groups according to their Hi-C contact intensities with the existing 90 groups. Haplotype phasing was performed in multiple rounds until no more contigs could be resolved. Chromosome assignment for each contig was based on genetic markers and alignments to the consensus assembly.

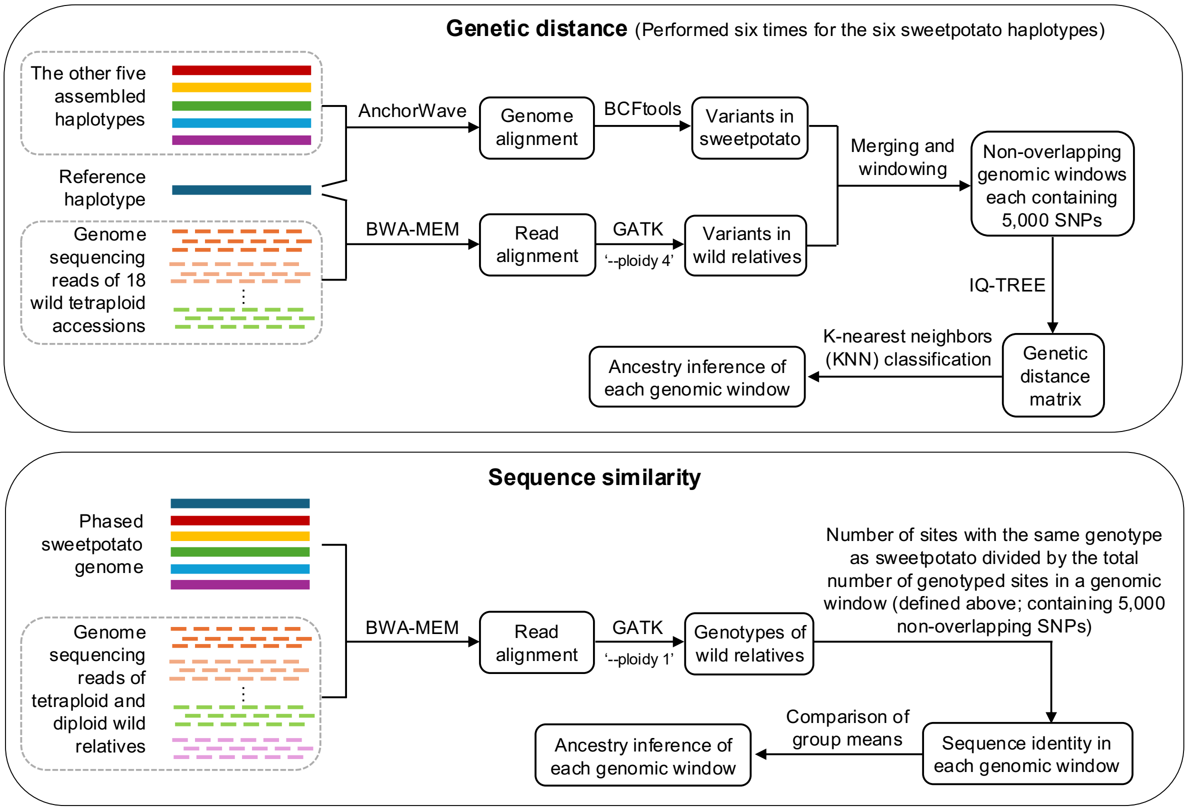


**Supplementary Figure 22** Ancestry inference of 'Tanzania'. Variants identified in sweetpotato haplotypes and wild accessions were used in phylogenetic analysis to generate a genetic distance matrix. Non-overlapping windows along the 'Tanzania' genome, each containing 5,000 SNPs, were classified into either *I. aequatoriensis* or *I. batatas* 4× type using a k-nearest neighbor (KNN) algorithm with seven nearest wild accessions of each sweetpotato haplotype. Sequence identity in each window between a wild relative and a sweetpotato haplotype was calculated as the number of sites with the same genotypes divided by the total number of genotyped sites in the window. Ancestry inference was based on a significant higher mean sequence identity for one group compared the other.
